# The cloud forest in the Dominican Republic: diversity and conservation status

**DOI:** 10.1101/543892

**Authors:** Ana Cano Ortiz, Carmelo M. Musarella, Ricardo Quinto Canas, José C. Piñar Fuentes, Carlos J. Pinto Gomes, Eusebio Cano

## Abstract

The study of the forest in rainy environments of the Dominican Republic reveals the presence of four types of vegetation formations, clearly differentiated from each other in terms of their floristic and biogeographical composition, and also significantly different from the rainforests of Cuba. This leads us to propose two new alliances and four plant associations located in northern mountain areas exposed to moisture-laden winds from the Atlantic: All. *Rondeletio ochraceae-Clusion roseae* (Ass. *Cyatheo furfuracei-Prestoetum motanae*; Ass. *Ormosio krugii-Prestoetum montanae*); and All. *Rondeletio ochraceae-Didymopanion tremuli* (Ass. *Hyeronimo montanae-Magnolietum pallescentis*; *Hyeronimo dominguensis-Magnolietum hamorii*). Due to human activity, some areas are very poorly conserved, as evidenced by the diversity index and the presence of endemic tree and plant elements. The worst conserved in terms of the relationship between characteristic plants vegetation (cloud forest) in areas with high rainfall in the Dominican Republic, along with its floristic diversity and state of conservation. Thanks to this study it has been possible to significantly increase the botanical knowledge of this important habitat.

## Introduction

The territory of the Dominican Republic (DR), with an extension of 48,198 km^2^ including the small adjacent islands, accounts for over two thirds of the territory of Hispaniola, an island located between parallels 17-19°N in the group of the Greater Antilles. Most previous botanical studies have concentrated predominantly on the flora –for example the work of [1] in the Sierra de Bahoruco– and highlight the abundant rainfall of up to 4,000 mm and the very high rate of endemic species. There are also other studies by several authors on the cloud forest in the Cordillera Central, Septentrional and Oriental ranges [2–17]. All these works, together with previous studies carried out by ourselves [18–29] have enabled us to undertake the present work. All the aforementioned studies focus attention on the knowledge of the flora, with only passing references to the vegetation. The main aim of this work is to determine the forest vegetation (cloud forest) in areas with high rainfall in the Dominican Republic, along with its floristic diversity and conservation status.

## Material and methods

The island of Hispaniola, with an area of 76,484 km^2^, and Cuba, Jamaica and Puerto Rico are the largest islands in the Caribbean region. The geological origin of the mountains on the island dates from the Cretaceous and Oligocene-Miocene era with the exception of the intramountain valleys formed during the Quaternary period due to the deposit of materials [30]. There is a predominance of calcareous materials with a karstic character, marbles, limestones and Quaternary deposit materials, and a large central nucleus of siliceous materials with serpentine outcrops [19–21]. The island has a mountainous relief with several mountain chains such as the Oriental, Central and Septentrional ranges, and sierras such as Bahoruco and Niebla. The northwest-southwest orientation of the mountains and the prevailing direction of the Atlantic winds explains the existence of a permanent sea of clouds, which gives rise to high rainfall on north-northeast-facing slopes.

This study is focused on the humid-hyper-humid forests in the Dominican Republic (DR) on the island of Hispaniola. Vegetation samples were taken in areas of high rainfall such as the Cordillera Central and Oriental ranges and the Sierra de Bahoruco, selecting sampling plots with an area of 500-2000 m^2^. Due to the scarcity of vegetation studies, we analysed the works of [31–35] in territories of Cuba. For the dynamic-catenal landscape study we took into account the criteria of [36–37]. An Excel© table was created with 483 rows (species) x 12 columns (tables containing 67 relevés) (Table 1). A statistical treatment (clustering) was applied to separate the communities described for Cuba from those of Hispaniola. The statistical treatment was done by adapting the Van der Maarel conversion [38] and substituting the abundance-dominance indexes with synthetic indexes with the following equivalence: I = 3, II = 4, III = 5, IV = 6, V = 7. Once the indexes were converted, a cluster analysis was applied using the Jaccard distance marking the distance between the associations studied. After separating the forests in the Dominican Republic (DR) from those of Cuba based on the Jaccard distance, an Excel© table was created with the vegetation relevés from the DR, and a Euclidean distance cluster analysis and a DCA were applied to obtain the different types of forests present in the DR. A CCA was done to determine the influence of environmental factors (temperature and rainfall) on the distribution of these forests, followed by a study of the diversity and conservation status.

**Table 1.** Plant communities studied and number of relevés.

|  | Communities | Authors | No. of rel. |
| --- | --- | --- | --- |
| C1 | *Clethro-Pinetum maestrensis* Borhidi 1991 (Cuba) | Borhidi (1991).<br>Phytogeography and vegetation ecology of Cuba, Table 139, page 624 | 5 |
| C2 | *Hyeronimo-Sloanetum curatellifoliae* Borhidi 1991 (Cuba) | Borhidi (1991).<br>Phytogeography and vegetation ecology of Cuba, Table 140, page 627 | 5 |
| C3 | *Alchorneo-Calophylletum rivularis* Reyes 2005 (Cuba) | Reyes (2005).<br>Synecological study of submontane rainforests on metamorphic complex rocks, Table 1 | 6 |
| C4 | *Pruno-Guareetum guidoniae* Reyes & Acosta 2011 (Cuba) | Reyes & Acosta (2011).<br>Phytocoenosis in evergreen forests in eastern Cuba III, Table 2 | 4 |
| C5 | *Ocoteo-Phoebietum elongatae* Reyes & Acosta 2010 (Cuba) | Reyes & Acosta (2010).<br>Phytocoenosis in evergreen forests in eastern Cuba I, Table 1 | 7 |
| C6 | *Guareo guidoniae-Zantoxyletum martinicensis* Reyes & Acosta 2010 (Cuba) | Reyes & Acosta (2010).<br>Phytocoenosis in evergreen forests in eastern Cuba II, Table 1 | 8 |
|  |  |  | Total 35 relevés |
| DR7 | *Hyeronimo montanae-Magnolietum pallescentis nova* (DR) | Own relevés | 5 |
| DR8 | *Cyatheo furfuracei-Prestoetum montanae nova* (DR) | Own relevés | 5 |
| DR9 | *Hyeronimo dominguensis-Magnolietum hamorii nova (DR)* | Own relevés | 4 |
| DR10 | *Ormosio krugii-Prestoetum montanae nova (DR)* | Own relevés | 3 |
| DR11 | Vegetation relevés (DR) | May & Peguero (2000). Vegetation and flora of the El Mogote ridge, Jarabacoa, Cordillera Central range, Dominican Republic; Table 1 page 23 | 3 |
| DR12 | Vegetation relevés (DR) | May (2007). Composition, structure and diversity in broadleaved cloud forests in the Ebano Verde scientific reserve (Cordillera Central range, Dominican Republic); Table 1 page 171 | 12 |
|  |  |  | Total 32 relevés |

## Results

The results of the analysis of Jaccard distances (Fig 1) applied to six plant communities in Cuba and six in the DR, show that the six communities described in Cuba by [31–35], C1-C6 (Table 1, 35 relevés), can be broken down into the community C1 and the group G_1_ (C2,C3,C4,C5,C6). C1 is differentiated from the rest in terms of its floristic, structural and ecological composition, as this is a pinewood of *Pinus maestrensis* Bise growing in rainy environments but on highly oligotrophic soils, in common with the other communities in group G_1_, which is floristically significantly different from group G_2_. There are very significant floristic differences between Cuba and the DR (Tables 2 and 3), with 173 species present in the samplings in the DR but not in Cuba, whereas the samplings in Cuba reveal 139 plants that are absent from the DR. In group G_2,_ which contains 32 of our own relevés and those one of [9–10], (DR7, DR8, DR9, DR10, DR11, DR12), the communities DR7, DR11 and DR12 can be seen to form a group for the DR representing different types of forests; these formations are a series of plant communities in very rainy environments in the Dominican Republic (DR) located in the Sierra de Bahoruco and the Cordillera Central and Oriental ranges, with rainfall of over 2,000 mm. Group G_2_ is broken down into two subgroups of plant communities –DR7-DR11-DR12 and DR8-DR9-DR10– which is plausible, as the first three correspond to areas with acid substrates and rainy environments in the Cordillera Central range, whereas the second subgroup contains communities growing on different kinds of substrates and in hyper-humid environments. We therefore focus on the analysis of 17 of our own samplings to which we apply a Euclidean distance cluster analysis and an ordination analysis, both of which perfectly separate the sampling groups.

**Fig 1.**
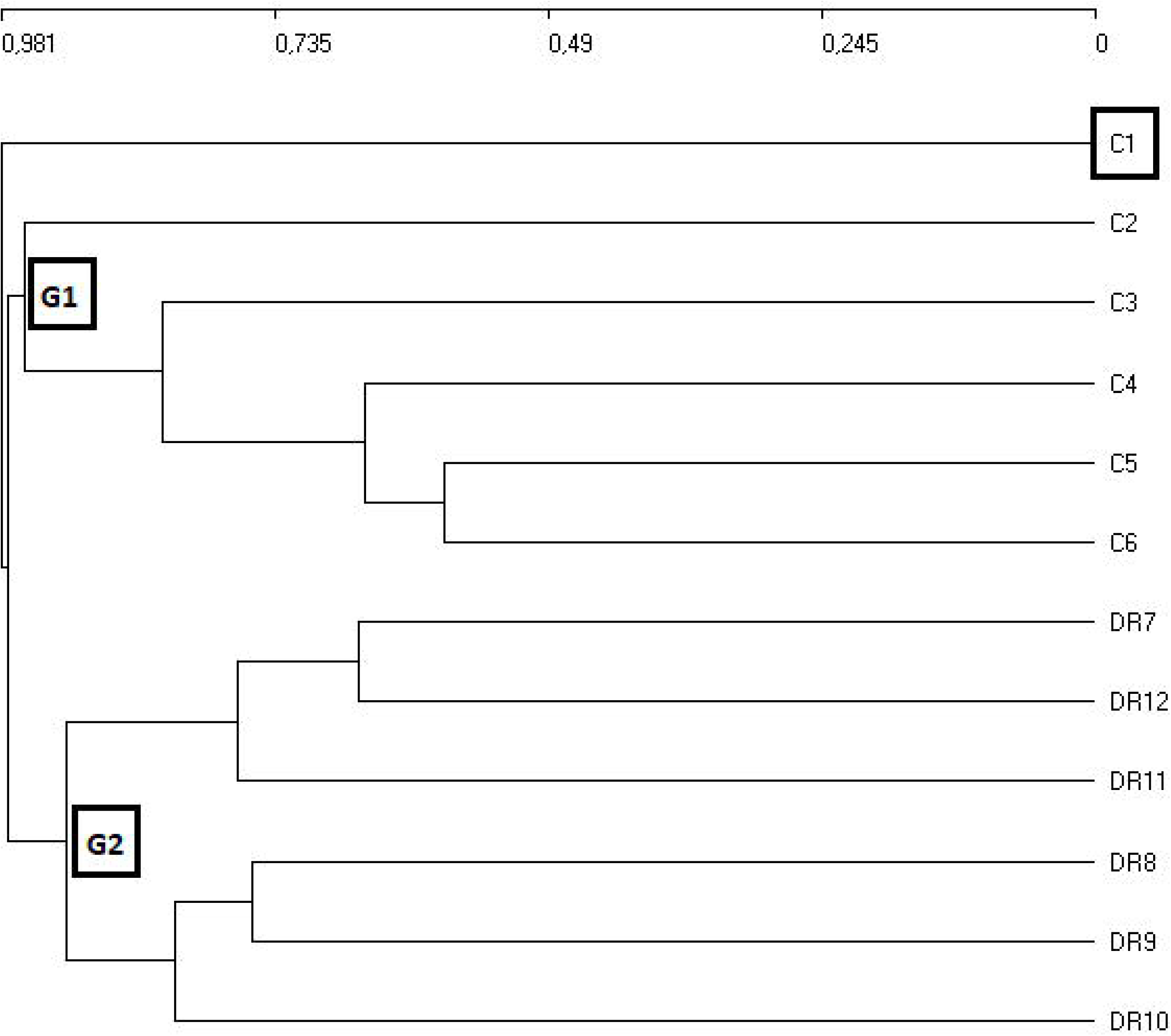
Jaccard distance cluster. Cluster analysis for the associations of Cuba and the Dominican Republic.

**Table 2.** Plants from Cuba not present in the relevés from the DR.

|  |  |
| --- | --- |
| *Ageratina paucibracteata* (Alain) King et Robins. | *Ocotea cuneata* (Griseb.) Urb. |
| *Alsophila aquilina* C. Chr. | *Oplismenus hirtellus* (L.) Beauv. |
| *Allophyllus cristalis* Lippold | *Ossaea muricata* (Griseb.) Wr. ex Sam. |
| *Ascyrum hypericoides* L. | *Ouratea revoluta* (Wr ex Griseb.) Engl. |
| *Bactris cubensis* Burret | *Panicum boliviense* Hack. |
| *Buchenavia capitata* (Vahl.) Eichl. | *Panicum glutinosum* Sw. |
| *Byrsonima biflora* Griseb. | *Panicum scoparium* L. |
| *Byrsonima coriacea* R. Br. | *Pardiaea maestrensis* Borhidi and Catassus |
| *Byrsonima orientensis* Bisse | *Pera ekmanii* Urb. |
| *Calophyllum utile* Bisse | *Phaius tankervilleae* (Banks) Blume |
| *Calycogonium grisebachii* Triana | *Pharus latifolius* L. |
| *Calycogonium lindenianum* Naud. | *Philodendron lacerum* (Jacq. ) Schott |
| *Calyptantes punctata* Griseb. | *Phyllanthus pachystylus* Urb. |
| *Callicarpa ferruginea* Sw. | *Phyllanthus subcarnosus* Wr ex Muell. Arg. |
| *Cestrum laurifolium* L'Hérit | *Pilocarpus racemosus* Vahl. |
| *Clethra cubensis* A. Rich. | *Pinus maestrensis* Bisse |
| *Clidemia strigillosa* (Sw.) DC. | *Piper holguinianum* Trel. |
| *Clusia minor* L. | *Pithecellobium arboreum* (L.) Urb. |
| *Clusia tetrastigma* Vesque | *Platyginia triandra* Borhidi |
| *Coccocypselum x lanceolatum* (Urb.) Borhidi | *Pleurothallis tribuloides* (Sw.) Lindl. |
| *Coccoloba costata* Wr. Inn Sauv. | *Pleurothallis velaticaulis* Rchb. |
| *Coccoloba ekmanii* O. C. Schmidt | *Podocarpus ekmanii* Urb. |
| *Coccoloba retusa* Griseb. | *Polygala oblongata* (Britt. ) Blake |
| *Coussarea urbaniana* Standl. | *Polypodium phyllitidis* L. |
| *Critonia dalea* (L.) DC. | *Polystachya cubensis* Schltr. |
| *Cyathea araneosa* (Sw.) Domin | *Protium cubense* (Rose) Urb. |
| *Cyrtilla nipensis* Urb. | *Psidium baliu* Urb. |
| *Chrysophyllum argenteum* Jacq. | *Psychotria shaferi* Urb. |
| *Dalbergaria cubensis* (Urb.) Borhidi | *Pteridium caudatum* (L.) Max. |
| *Dendropanax nervosus* (Urb et Ekm.) A. C. Sm. | *Pteris rigida* Sw. |
| *Dennstaedtia adiantoides* (H. & B) Moore | *Pytirogramma sulphurea* (Sw.) Max. |
| *Desmodium herbaceum* (L.) Benth. & Oerst. | *Rajania nipensis* Howard |
| *Dicranopteris flexuosa* (Schrad.) Mett. | *Raudolfia salicifolia* Griseb. |
| *Dichaea hystericina* Rchb. | *Rhamnidium nipense* Urb. |
| *Dilomilis oligophylla* (Schltr.) Summerh. | *Rondeletia calophylla* Standl ex Britt. |
| *Dinema cubincola* (Borhidi) H. Dietr. | *Rondeletia naguensis* Britt and Wils |
| *Dipholis cubensis* (Griseb.) Pierre | *Rubus turquinensis* Rydb. |
| *Ditta myricoides* Griseb. | *Salacia nipensis* Britt. |
| *Elaphoglossum firmum* (Mett.) Urb. | *Sapium cubense* Britt ex Wils. |
| *Eugenia rigida* Berg. | *Sapium erythrospermum* (Griseb.) Muell. Arg. |
| *Garrya fadyenii* Hook. | *Sapium jamaicense* Sw. |
| *Gesneria pachyclada* Urb. | *Savia cuneifolia* Urb. |
| *Gesneria viridiflora* (Dcne) Kuntze | *Scleria pilosissima* Britt. |
| *Graffenrieda refescens* Britt. & Wils. | *Schradera cubensis* Steyerem. |
| *Guatteria moralesii* Urb. | *Simaruba laevis* Griseb. |
| *Guettarda lindeniana* A. Rich. | *Sloanea curatellifolia* Griseb. |
| *Habenaria monorrhiza* (Sw.) Rchb. | *Solanum cristalense* Amsh. |
| *Hedyosmum grisebachii* Solms | *Solanum pachyneurum* O.E. Schulz |
| *Heterotrichum umbellatum* (Mill.) Urb. | *Solanum virgatum* Lam. |
| *Hyeronima nipensis* Urb. | *Solonia reflexa* Urb. |
| *Ichnanthus mayarensis* (Wr.) Hitchc. | *Stelis ophioglossoides* (Jacq.) Sw. |
| *Illicium cubense* A.C.Sm. | *Tabebuia hypoleuca* Griseb. |
| *Jacquiella globosa* (Jacq.) Schlechter | *Talauna minor* Urb. subsp. *oblongifolia* (León) Borhidi |
| *Lasiacis sloanei* (L.) Hitchc. | *Terminalia nipensis* Alain |
| *Leucocroton wrightii* Griseb. | *Trema cubensis* Urb. |
| *Litachne pauciflora* (Swartz) P. Beauverd | *Trichomanes crispum* L. |
| *Lobelia assurgens* L. | *Vaccinium leonis* Acuña & Roig |
| *Lophosoria quadripinnata* (Gmel.) C. Chr. | *Vandenboschia scandens* (L.) Copel |
| *Lycopodium dichotomum* Jacq. | *Vanilla phaeantha* Rchb. |
| *Lyonia calycosa* (Samll) Urb. | *Vanilla wrightii* Rchb. |
| *Lyonia maestrensis* Acuña and Roig | *Vernonia parvuliceps* Ekm. |
| *Magnolia cubensis* Urb. | *Viburnum villosum* Sw. |
| *Marcgravia evenia* Kr et Urb. | *Wallenia laurifolia* (A. Rich.) Wr in Sauv subsp. *pinetorum* (Britt.) Borhidi |
| *Mataiba domingensis* (DC.) Radlk. | *Wigandia reflexa* Brand |
| *Mecodium polyanthos* (Sw.) Copel | *Zanthoxylum cubense* P. Wils. |
| *Meriania leucantha* Sw. subsp. *nana* (Naud.) Borhidi |  |
| *Meringium fucoides* (Sw.) Copel |  |
| *Miconia acunae* Borhidi |  |
| *Miconia alternifolia* Griseb. |  |
| *Miconia dodecandra* (Desv.) Cogn. |  |
| *Micropholis polita* (Griseb.) Pierre |  |
| *Mozartia gundlachii* (Kr. & Urb.) Urb. |  |
| *Myrica cacuminis* Britt and Wils. |  |
| *Myrica punctata* Griseb. |  |

**Table 3.** Plants from the Dominican Republic (DR) not present in the relevés from Cuba.

|  |  |
| --- | --- |
| *Alsophila minor* (D.C.Eaton) R.M.Tryon | *Magnolia hamorii* Howard |
| *Anacheilium cochleatum* (L.) Hoffm. | *Magnolia pallescens* Urb. & Ekm. |
| *Antrophyum lanceolatum* (L.) Kaulf. | *Malpighia macracantha* Ekm. & Nied. |
| *Arthrostylidium multispicatum* Pilger | *Marattia kaulfussii* J. Smith |
| *Arthrostylidium sarmentosum* Pilger | *Marcgravia rubra* A. Liogier |
| *Asplenium radicans* L. | *Maxillaria coccinea* (Jacq.) L.O. Wms. |
| *Baccharis myrsinites* (Lam.) Pers. | *Mecranium ovatum* Cog. |
| *Bactris plumeriana* Mart. | *Meriania involucrata* (Desv.) Naud. |
| *Beilschmiedia pendula* (Sw.) Hemsl. | *Miconia mirabilis* (Aubl.) L.O. Willians |
| *Blechnum fragile* (Liebm.) Morton & Lellinger | *Miconia prasina* (Sw.) DC. |
| *Blechnum tuerckheimii* A. Brause | *Miconia racemosa* (Aubl.) DC. |
| *Bocconia frutescens* L. | *Microgramma piloselloides* L. |
| *Buchenavia tetraphylla* (Aubl.) R. A. Howard | *Mikania cordifolia* (L.) Willd. |
| *Byrsonima lucida* (Mill.) L.c. rich. | *Mikania venosa* A. Liogier |
| *Byrsonima spicata* (Cav.) Kunth | *Mimosa pudica* L. |
| *Calyptantes selleanus* Urb. & Ekm. | *Mucuna urens* (L.) Fawc. & Rendle |
| *Camparettia falcata* Poepp. & Endl. | *Myrcia deflexa* (Poir) DC. |
| *Casearea arborea* (L.C.Rich.) Urb. | *Myrsine nubicola* A. Liogier |
| *Cecropia screberiana* Miq. | *Nephrolepis multiflora* (Roxb.) Jarret |
| *Cestrum coelophlebium* O. E. Schulz | *Nepsera aquatica* (Aubl.) Naud. |
| *Cestrum daphnoides* Griseb. | *Neurolaena lobata* (L.) Cass. |
| *Cestrum inclusum* Urb. | *Niphidium crassifolium* (L.) Lell. |
| *Cinnamomum alainii* (C.K. Allen) A. Liogier | *Ocotea acarina* C. K. allen |
| *Cissampelos pareira* L. | *Ocotea floribunda* (Sw.) Mez |
| *Cissus verticillata* (L.) Nicholson & Farris | *Ocotea foeniculacea* Mez |
| *Clidemia umbellata* (Miller) L.O. Wms. | *Ocotea globosa* (Aubl.) Schlecht. & Cham. |
| *Clusia clusioides* (Griseb.) D'arcy | *Ocotea nemodaphne* Mez |
| *Cnemidaria horrida* (L.) K. Presl | *Ocotea patens* (Sw.) Nees |
| *Coccoloba wrightii* Lindau | *Odontadenia polyneura* (Urb.) Wood. |
| *Columnea domingensis* (Urb.) Wiehler | *Olyra latifolia* L. |
| *Columnea sanguinea* Urb. | *Oncidium variegatum* (Sw.) Sw. |
| *Commelina elegans* Kunth | *Ophioglossum palmatum* L. |
| *Cordia dependens* Urb. & Ekm. | *Oreopanax capitatus* (Jacq.) Decne. & Planch. |
| *Cupania americana* L. | *Ormosia krugii* Urb. |
| *Cyathea fulgens* C. Chr. | *Palicourea crocea* (Sw.) Schultes |
| *Cyathea furfuracea* Baker | *Passiflora rubra* L. |
| *Chaetocarpus domingensis* Proctor | *Peperomia hernandifolia* (Vahl) A. Dietr. |
| *Daphnopsis crassifolia* (Poir.) Meiss. | *Persea krugii* Mez |
| *Dendropanax arboreus* (L.) Dcne & Planch. | *Persea oblongifolia* Kopp. |
| *Dichaea glauca* (Sw.) Lindley | *Phlebodium aureum* (L.) J. Smith |
| *Didymopanax tremulus* Krug. & Urb. | *Pilea geminata* Urb. |
| *Dilomilis montana* (Sw.) Summerh. | *Pinguicula casabitoana* J. Jiménez |
| *Diplazium hastile* (Christ.) C. Chr. | *Piper adunculum* L. |
| *Diplazium hians* Kuntze | *Pleurothallis ruscifolia* (Jacq.) R. Br. |
| *Ditta maestrensis* Borhidi | *Pleurothallis domingensis* Cogn. |
| *Elaphoglossum crinitum* (L.) C. Chr. | *Polygala fuertesii* (Urb.) Blake |
| *Elaphoglossum latifolium* (Sw.) J. Sm. | *Polypodium angustifolium* Sw. |
| *Elleanthus cephalotus* Garay & Sweet | *Polypodium loriceum* L. |
| *Entada gigas* (L.) Fawc. & Rendle | *Pothomorphe peltata* (L.) Miquel |
| *Epidendrum anceps* Jacq. | *Pothuya nudicaulis* (L.) Regel |
| *Epidendrum carpophorum* Barb. Rodr. | *Prestoea montana* (Grah.) Nichol |
| *Epidendrum jamaicense* Lindl | *Psychotria domingensis* Jacq. |
| *Epidendrum ramosum* Jacq. | *Psychotria liogieri* Sateyerm |
| *Eupatorium odoratum* L. | *Psychotria uliginosa* Sw. |
| *Exostema elliptica* Griseb. | *Pytirogramma calomelanos* (L.) Link |
| *Gleychenia bifida* (Willd.) Spreng. | *Renealmia jamaicensis* (Gaertn.) Horan var. *puberula* (Gagn.) Maas |
| *Gomedesia lindeniana* Berg. | *Rondeletia ochracea* Urb. |
| *Gonocalyx tetrapteris* A. Liogier | *Sagraea fuertesii* (Cogn.in Urb.) Alain |
| *Grammitis asplenifolia* (L.) Proctor | *Schlegelia brachyantha* Griseb. |
| *Guarea guidonea* Sleumer | *Schradera subsessilis* Steyermark |
| *Guatteria blainii* (Griseb.) Urb. | *Senecio lucens* (Poir.) Urb. |
| *Guzmania monostrachya* (Sw.) Rusby | *Sloanea berteriana* Choisy |
| *Gyrotaenia myriocarpa* Griseb. | *Smilax havanensis* Jacq. |
| *Hedychium coronarium* Koen. | *Smilax populnea* Kunt var. *horrida* O.E. Schulz |
| *Hedyosmum domingense* Urb. | *Solanum crotonoides* Lam. |
| *Hirtella triandra* Sw. | *Solanum jamaicense* Mill. |
| *Hyeronima domingensis* Urb. | *Solanum torvum* Sw. |
| *Hyeronima montana* A. Liogier | *Solanum virgatum* Lam. |
| *Hypolepis hispaniolica* Mason | *Stigmaphyllon emarginatum* (L.) A. Juss. |
| *Hyptis americana* (Poir.) Briq. | *Styrax ochraceus* Urb. |
| *Ichnanthus pallens* (Sw.) Munro | *Syngonium podophyllum* Schott |
| *Ilex tuerckheimii* Loes. | *Tabebuia bullata* A. Gentry |
| *Inga fagifolia* (L.) Willd. ex Benth. | *Tabebuia vinosa* A. Gentry |
| *Inga vera* Willd. | *Torrallbasia cuneifolia* (C. Wright) Krug. & Urb. |
| *Ipomoea furcyensis* Urb. | *Triunfetta semitriloba* Jacq. |
| *Ipomoea tiliacea* (Willd.) Choisy | *Turpinia occidentalis* (Sw.) G. Don |
| *Isachne rigidifolia* (Poir.) Urb. | *Uncinia hamata* (L.) Urb. |
| *Lasianthus bahorucanus* Zannoni | *Urena lobata* L. |
| *Leandra limoides* (Urb.) W. Judd & Skean | *Urera baccifera* (L.) Gaud. |
| *Lobelia robusta* Graham | *Vaccinium racemosum* (Vahl) Wilbur & Luteyn |
| *Lobelia rotundifolia* Juss. | *Vernonia buxifolia* (Cass.) Less. |
| *Lomariopsis sorbifolia* (L.) Fée | *Vitis tiliifolia* H. & B. ex Willd. |
| *Lycopodium cernuum* L. | *Vittaria lineata* (L.) Smith |
| *Lycopodium clavatum* L. | *Vriesea sintenisii* (Baker) L.B. Smith & Pitt. |
| *Lyonia alainii* W. Judd. | *Vriesea tuercheimii* (Mez.) L.B. Smith |
| *Macrocarpaea domingensis* Urb. | *Zanthoxylum martinicensis* (DC.) Lam. |
| *Machaerina cubensis* (Kük.) T. Koyama |  |

### Phytosociological study

The statistical analysis of the samplings from the DR reveals the existence of four forest plant associations (Fig 2): As1) *Hyeronimo montanae-Magnolietum pallescentis nova hoc loco* (S1 Table rel. DR1, DR2, DR4, DR5, DR6; typus rel. DR4), growing at altitudes of between 1,300 and 1,500 metres on siliceous substrates in the Cordillera Central range (central biogeographical district), and in rainy environments with a humid ombrotype and a mesotropical thermotype [16,23,39,40]. These forests contact in hyper-humid areas with forests of *Prestoea montana* (Grah.) Nichol, and have a high floristic diversity with 21 trees, eight climbing species and five epiphytes, and a high rate of endemisms (14 species). As2) *Cyatheo furfuracei-Prestoetum montanae nova hoc loco* (S2 Table rel. DR3, DR7, DR8, DR9, DR10; typus rel. DR3), a plant community dominated by *Prestoea montana*, always found in hyper-humid environments, generally in very rainy and shady gorges, contacting with the previous association towards areas that are somewhat less rainy and more exposed to sun and wind. It also has a high diversity, with 40 tree and 25 epiphyte species. Due to the catenal contact between both associations, As1 and As2 present a series of common species; they are therefore statistically close (Figs 3 and 4). As3) *Hyeronimo dominguensis-Magnolietum hamorii nova hoc loco* (S3 Table rel. DR11, DR12, DR13, DR14; typus rel. DR11) represents forests of *Magnolia* in the Sierra de Bahoruco, which develop on calcareous substrates in humid environments at altitudes of around 1,200-1,300 metres in a humid ombrotype and a mesotropical thermotype, with a high number of tree (25) and epiphyte (14) species. As4) *Ormosio krugii-Prestoetum montanae nova hoc loco* (S4 Table rel. DR15, DR16, DR17; typus rel. DR16), an association characterised by a high diversity of trees (27 species), and a lower number of endemic species than the previous associations. The four associations present a clear floristic and biogeographical differentiation (Fig 5, Table 4) [41,42].

**Fig 2.**
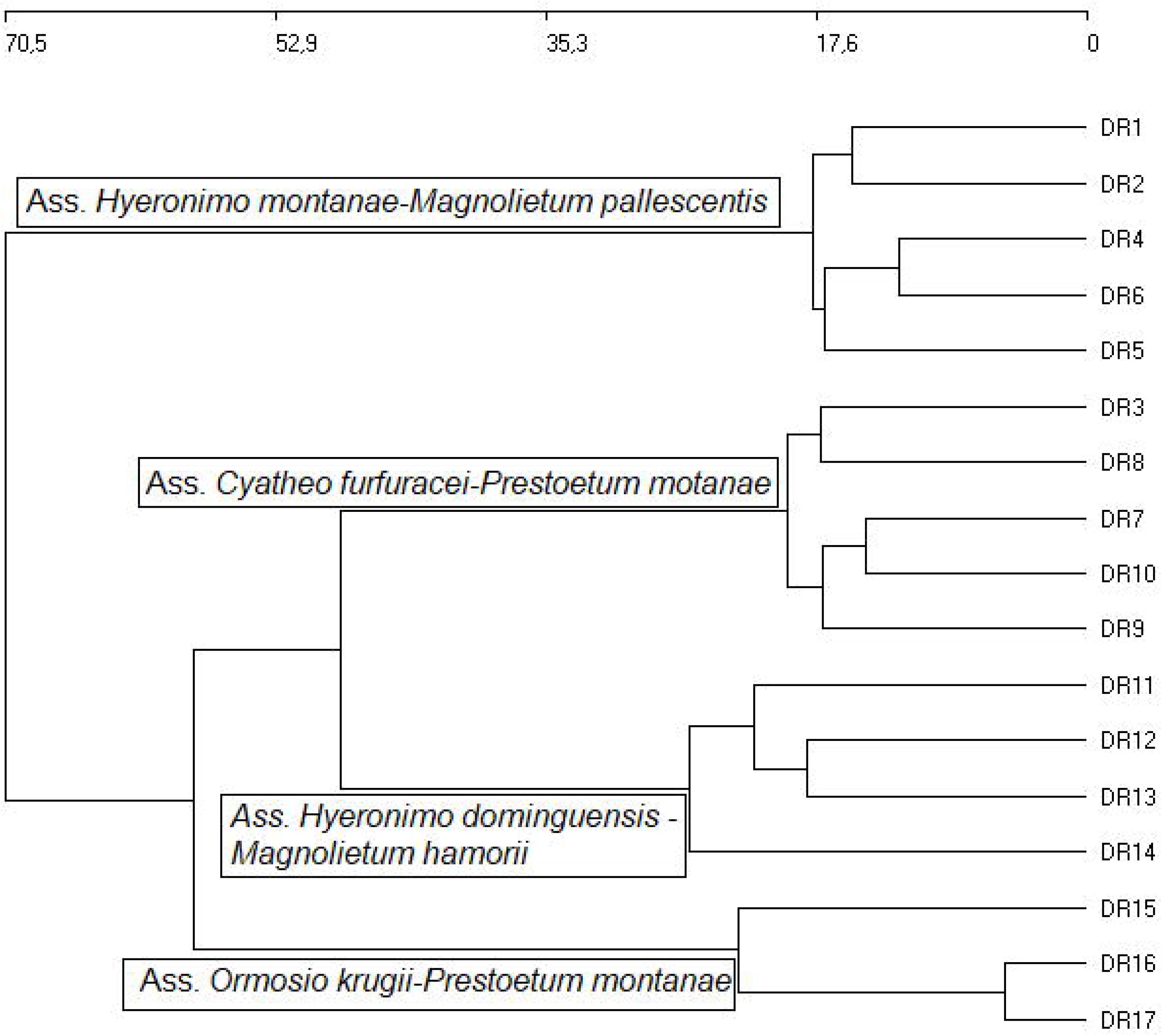
Cluster from the DR. Euclidean distance using Ward’s method.

**Fig 3.**
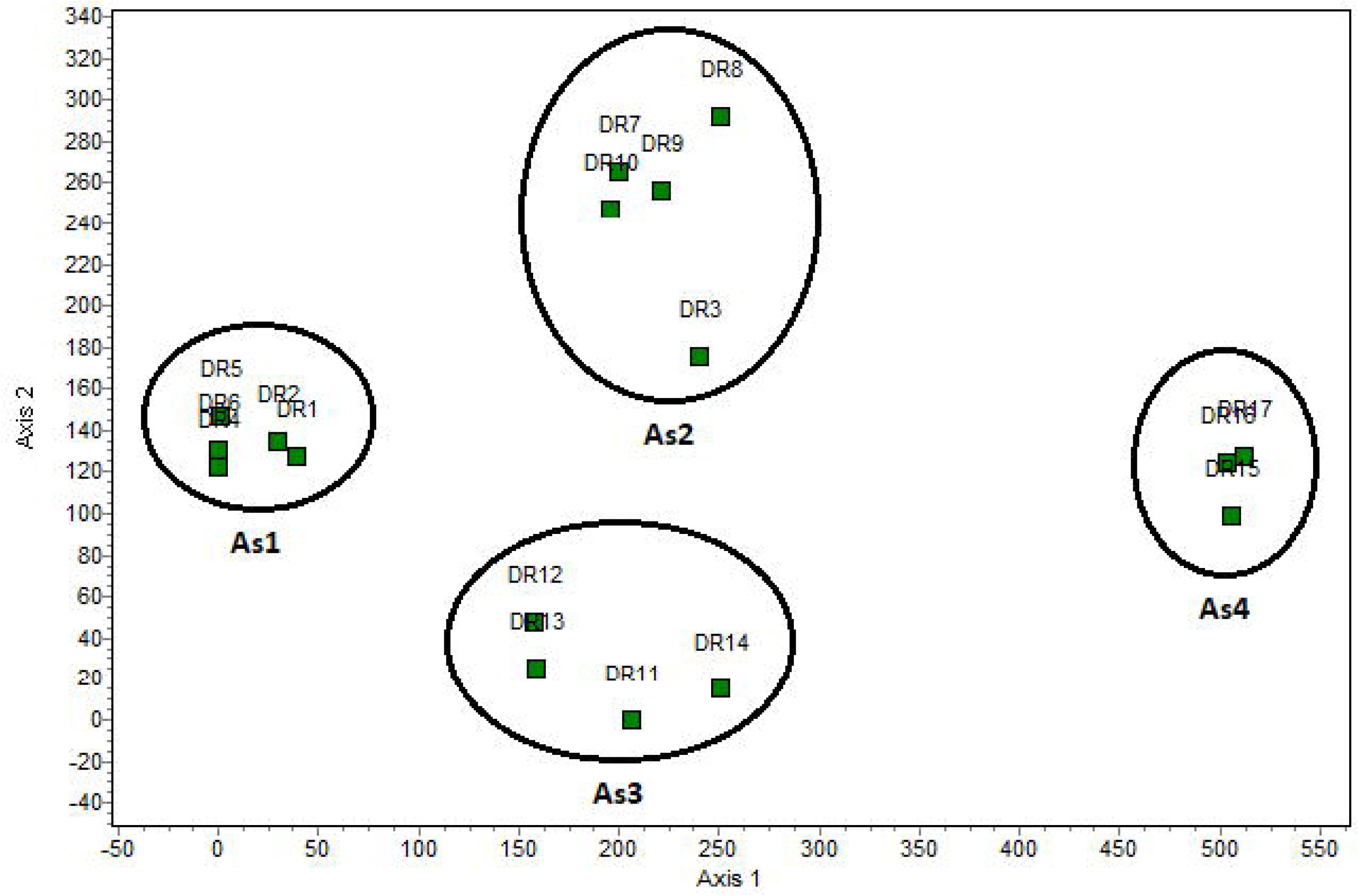
DCA ordination analysis. Management analysis for inventories of the Dominican Republic, separation between 4 associations.

**Fig 4.**
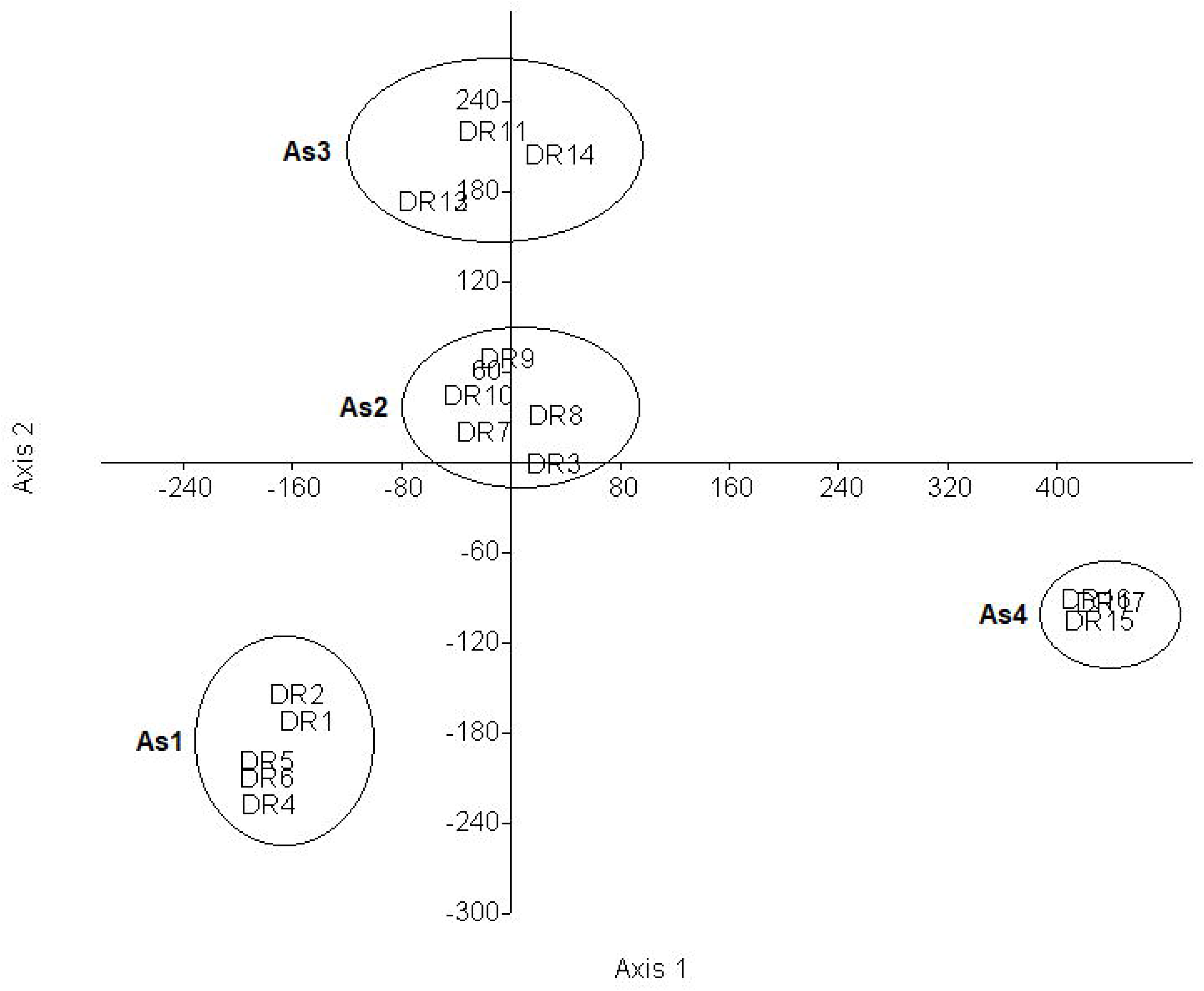
DCA ordination analysis. DCA analysis confirming the separation of the 4 associations.

**Fig 5.**
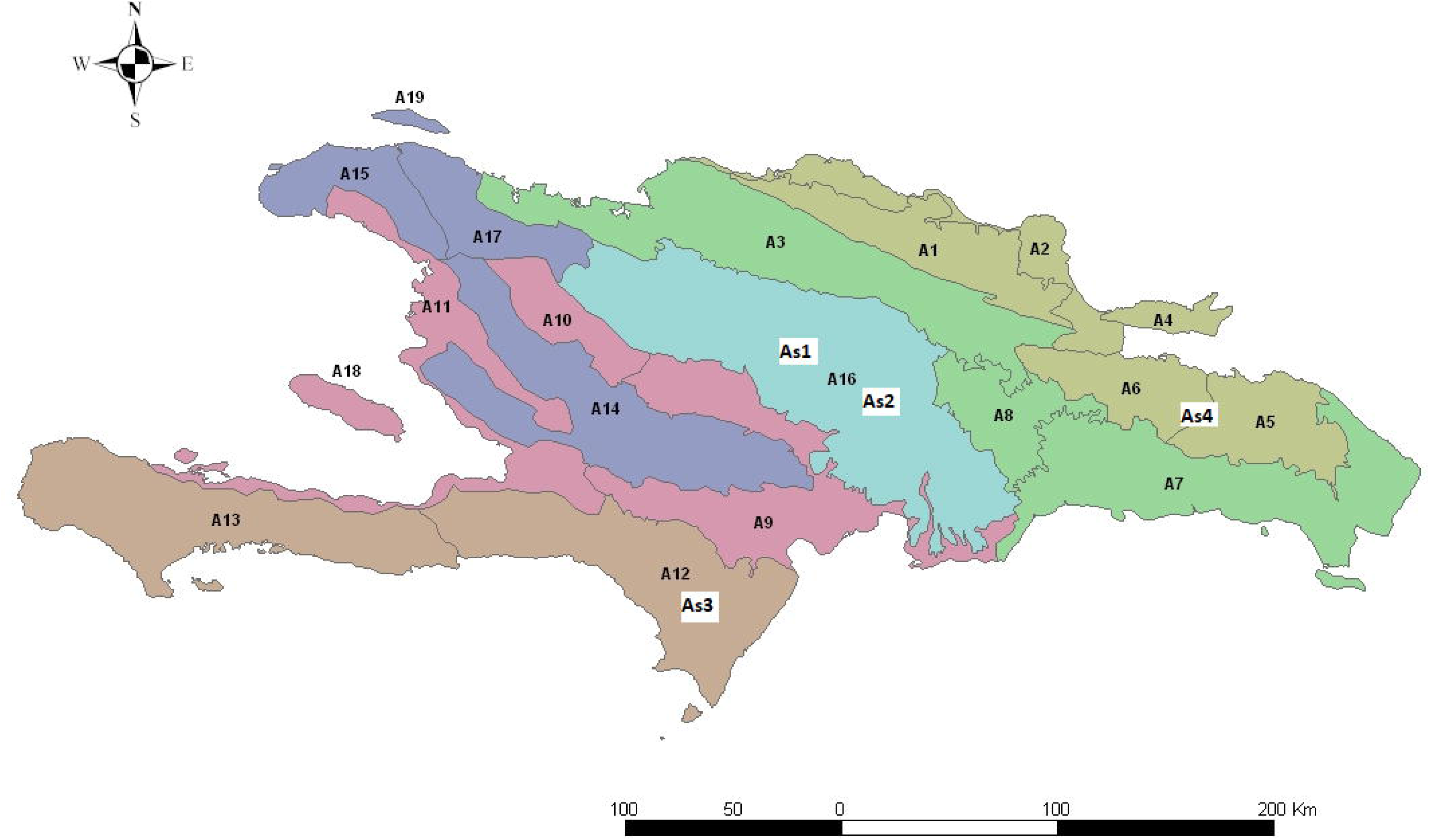
Biogeographical distribution of the associations in the study. As1. *Hyeronimo montanae-Magnolietum pallescentis* (A16: central district). As2. *Cyatheo furfuracei-Prestoetum montanae* (A16: central district). As3. *Hyeronimo dominguensis-Magnolietum hamorii* (A12: Bahoruco district). As4. *Ormosio krugii-Prestoetum montanae* (A5: eastern district).

**Table 4.** Synthetic table of the associations in the study.

| Synthetic table | As1 | As2 | As3 | As4 | P |
| --- | --- | --- | --- | --- | --- |
| *Myrsine coriacea* (Sw.) R. Br. | IV | III | V | III | 4 |
| *Ocotea leucoxylon* (Sw.) Mez | I | IV | III | III | 4 |
| *Prestoea montana* (Grah.) Nichol | I | V | V | V | 4 |
| *Psychotria domingensis* Jacq. | IV | IV | V | III | 4 |
| *Gleychenia bifida* (Willd.) Spreng. | II | I |  | I | 3 |
| *Clidemia umbellata* (Miller) L.O. Wms. | I | I | - | I | 3 |
| *Renealmia jamaicensis* (Gaertn.) Horan var. *puberula* (Gagn.) Maas | V | III | V | - | 3 |
| *Arthrostylidium multispicatum* Pilger | V | IV | III | - | 3 |
| *Rondeletia ochracea* Urb. | V | II | V | - | 3 |
| *Didymopanax tremulus* Krug. & Urb. | IV | I | V | - | 3 |
| *Psychotria guadalupensis* (DC.) Howard | III | III | V | - | 3 |
| *Mikania venosa* A. Liogier | II | IV | V | - | 3 |
| *Odontosoria uncinella* (Kunze) Fée | II | II | V | - | 3 |
| *Brunellia comocladifolia* H. & B. | II | III | III | - | 3 |
| *Lobelia rotundifolia* Juss. | III | I | I | - | 3 |
| *Alchornea latifolia* Sw. | - | III | I | V | 3 |
| *Miconia mirabilis* (Aubl.) L.O. Willians | - | II | I | V | 3 |
| *Mucuna urens* (L.) Fawc. & Rendle | - | II | I | V | 3 |
| *Nephrolepis multiflora* (Roxb.) Jarret | - | I | III | I | 3 |
| *Ilex macfadyenii* (Walp.) Rehder | V | - | I | - | 2 |
| *Chionanthus domingensis* Lam. | V | - | I | - | 2 |
| *Macrocarpaea domingensis* Urb. | IV | - | III | - | 2 |
| *Polygala fuertesii* (Urb.) Blake | IV | - | III | - | 2 |
| *Marcgravia rubra* A. Liogier | IV | - | I | - | 2 |
| *Alsophila minor* (D.C.Eaton) R.M.Tryon | V | III | - | - | 2 |
| *Palicourea alpina* (Sw.) DC. | V | II | - | - | 2 |
| *Blechnum occidentale* L. | III | III | - | - | 2 |
| *Cyrilla racemiflora* L. | IV | IV | - | - | 2 |
| *Ocotea nemodaphne* Mez | III | - | I | - | 2 |
| *Schradera subsessilis* Steyermark | II | - | I | - | 2 |
| *Lycopodium clavatum* L. | II | - | - | I | 2 |
| *Odontadenia polyneura* (urb.) Wood. | II | I | - | - | 2 |
| *Byrsonima lucida* (Mill.) DC. | I | II | - | - | 2 |
| *Weinmannia pinnata* L. | I | - | V | - | 2 |
| *Epidendrum carpophorum* Barb. Rodr. | I | - | I | - | 2 |
| *Epidendrum carpophorum* Barb. Rodr. | I | - | I | - | 2 |
| *Pleurothallis domingensis* Cogn. | I | II | - | - | 2 |
| *Cestrum coelophlebium* O. E. Schulz | I | II | - | - | 2 |
| *Olyra latifolia* L. | I | I | - | - | 2 |
| *Cecropia screeberiana* Miq. | - | III | - | V | 2 |
| *Turpinia occidentalis* (Sw.) G. Don | - | III | - | V | 2 |
| *Mikania cordifolia* (L.) Willd. | - | I | - | V | 2 |
| *Pothomorphe peltata* (L.) Miquel | - | I | - | III | 2 |
| *Ichnanthus pallens* (Sw.) Munro | - | III | - | I | 2 |
| *Guzmania monostrachya* (Sw.) Rusby | - | II | - | I | 2 |
| *Dendropanax arboreus* (L.) Dcne & Planch. | - | IV | I | - | 2 |
| *Dichaea glauca* (Sw.) Lindley | - | III | III | - | 2 |
| *Epidendrum ramosum* Jacq. | - | III | I | - | 2 |
| *Gomedesia lindeniana* Berg. | - | II | V | - | 2 |
| *Myrcia deflexa* (Poir) DC. | - | II | V | - | 2 |
| *Peperomia hernandifolia* (Vahl) A. Dietr. | - | II | V | - | 2 |
| *Vriesea tuercheimii* (Mez.) L.B. Smith | - | I | V | - | 2 |
| *Cyathea fulgens* C. Chr. | - | I | V | - | 2 |
| *Magnolia hamorii* Howard | - | I | V | - | 2 |
| *Mecranium ovatum* Cog. | - | I | V | - | 2 |
| *Lasianthus bahorucanus* Zanoni | - | I | V | - | 2 |
| *Nephrolepis biserrata* (Sw.) Schott | - | I | V | - | 2 |
| *Columnnea domingensis* (Urb.) Wiehler | - | I | V | - | 2 |
| *Hedyosmum domingense* Urb. | - | I | III | - | 2 |
| *Lomariposis sorbifolia* (L.) Feé | - | I | III | - | 2 |
| *Beilschmiedia pendula* (Sw.) Hemsl. | - | I | III | - | 2 |
| *Vaccinium racemosum* (Vahl) Wilbur & Luteyn | IV | I | III | - | 2 |
| *Ocotea acarina* C. | - | I | I | - | 2 |
| *Hypolepis hispaniolica* Mason | - | I | I | - | 2 |
| *Schlegelia brachyantha* Griseb. | - | I | I | - | 2 |
| *Sagraea fuertesii* (Cogn.in Urb.) Alain | - | II | I | - | 2 |
| *Niphidium crassifolium* (L.) Lell. | - | I | I | - | 2 |
| *Phlebodium aureum* (L.) J. Smith | - | I | I | - | 2 |
| *Polypodium loriceum* L. | - | I | I | - | 2 |
| *Epidendrum jamaicense* Lindl | - | I | I | - | 2 |
| *Microgramma piloselloides* L. | - | I | - | I | 2 |
| *Miconia prasina* (Sw.) DC. | - | - | III | V | 2 |
| *Guarea guidonea* Sleumer | - | - | I | V | 2 |
| *Tibouchina longifolia* (Vahl) Baill. | - | - | I | III | 2 |
| *Smilax domingensis* Willd. | - | - | I | I | 2 |
| *Magnolia pallescens* Urb. & Ekm. | V | - | - | - | 1 |
| *Styrax ochraceus* Urb. | V | - | - | - | 1 |
| *Hyeronima montana* A. Liogier | V | - | - | - | 1 |
| *Cyathea furfuracea* Baker | V | - | - | - | 1 |
| *Clusia clusioides* (Griseb.) D'arcy | V | - | - | - | 1 |
| *Ditta maestrensis* Borhidi | V | - | - | - | 1 |
| *Persea oblongifolia* Kopp. | V | - | - | - | 1 |
| *Smilax populnea* Kunt var. *horrida* O.E. Schulz | V | - | - | - | 1 |
| *Tabebuia vinosa* A. Gentry | V | - | - | - | 1 |
| *Gonocalyx tetrapterus* A. Liogier | V | - | - | - | 1 |
| *Cinnamomum alainii* (C.K. Allen) A. Liogier | IV | - | - | - | 1 |
| *Vriesea sintenisii* (Baker) L.B. Smith & Pitt. | III | - | - | - | 1 |
| *Baccharis myrsinites* (Lam.) Pers. | III | - | - | - | 1 |
| *Pinguicula casabitoana* J. Jiménez | III | - | - | - | 1 |
| *Chaetocarpus domingensis* Proctor | II | - | - | - | 1 |
| *Odontosoria aculeata* (L.) J. Sm. | I | - | - | - | 1 |
| *Myrsine nubicola* A. Liogier | I | - | - | - | 1 |
| *Persea krugii* Mez | I | - | - | - | 1 |
| *Lycopodium cernuum* L. | I | - | - | - | 1 |
| *Isachne rigidifolia* (Poir.) Urb. | I | - | - | - | 1 |
| *Machaerina cubensis* (Kük.) T. Koyama | I | - | - | - | 1 |
| *Vernonia buxifolia* (Cass.) Less. | I | - | - | - | 1 |
| *Lyonia alainii* W. Judd. | I | - | - | - | 1 |
| *Clidemia hirta* (L.) D. don | I | - | - | - | 1 |
| *Bocconia frutescens* L. | I | - | - | - | 1 |
| *Dilomilis montana* (Sw.) Summerh. | I | - | - | - | 1 |
| *Myrcia splendens* (Sw.) DC. | - | IV | - | - | 1 |
| *Cissampelos pareira* L. | - | III | - | - | 1 |
| *Uncinia hamata* (L.) Urb. | - | III | - | - | 1 |
| *Tabebuia bullata* A. Gentry | - | III | - | - | 1 |
| *Blechnum tuerckheimii* A. Brause | - | III | - | - | 1 |
| *Senecio lucens* (Poir) Urb. | - | III | - | - | 1 |
| *Coccoloba wrightii* Lindau | - | III | - | - | 1 |
| *Guatteria blainii* (Griseb.) Urb. | - | II | - | - | 1 |
| *Solanum crotonoides* Lam. | - | II | - | - | 1 |
| *Vitis tiliifolia* H. & B. ex Willd. | - | I | - | - | 1 |
| *Anacheilium cochleatum* (L.) Hoffm. | - | I | - | - | 1 |
| *Antrophyum lanceolatum* (L.) Kaulf. | - | I | - | - | 1 |
| *Camparettia falcata* Poepp. & Endl. | - | I | - | - | 1 |
| *Passiflora rubra* L. | - | I | - | - | 1 |
| *Smilax havanensis* Jacq. | - | I | - | - | 1 |
| *Stigmaphyllon emarginatum* (L.) A. Juss. | - | I | - | - | 1 |
| *Commelina elegans* Kunth | - | I | - | - | 1 |
| *Diplazium hastile* (Christ.) C. Chr. | - | I | - | - | 1 |
| *Diplazium hians* Kuntze | - | I | - | - | 1 |
| *Epidendrum anceps* Jacq. | - | I | - | - | 1 |
| *Grammitis asplenifolia* (L.) Proctor | - | I | - | - | 1 |
| *Jacquinella globosa* (Jacq.) Schlechter | - | I | - | - | 1 |
| *Oncidium variegatum* (Sw.) Sw. | - | I | - | - | 1 |
| *Pothuya nudicaulis* (L.) Regel | - | I | - | - | 1 |
| *Vittaria lineata* (L.) Smith | - | I | - | - | 1 |
| *Cestrum inclusum* Urb. | - | I | - | - | 1 |
| *Cordia dependens* Urb. & Ekm. | - | I | - | - | 1 |
| *Daphnopsis crassifolia* (Poir.) Meiss. | - | I | - | - | 1 |
| *Eupatorium odoratum* L. | - | I | - | - | 1 |
| *Gyrotaenia myriocarpa* Griseb. | - | I | - | - | 1 |
| *Hyptis americana* (Poir.) Briq. | - | I | - | - | 1 |
| *Lasianthus lanceolatus* (Griseb.) Gómez Maza |  | I | - | - | 1 |
| *Lobelia robusta* Graham | - | I | - | - | 1 |
| *Psychotria liogieri* Sateyerm | - | I | - | - | 1 |
| *Solanum virgatum* Lam. | - | I | - | - | 1 |
| *Pilea geminata* Urb. | - | I | - | - | 1 |
| *Exostema elliptica* Griseb. | - | I | - | - | 1 |
| *Malpighia macracantha* Ekm. & Nied. | - | I | - | - | 1 |
| *Ocotea floribunda* (Sw.) Mez | - | I | - | - | 1 |
| *Ocotea patens* (Sw.) Nees | - | I | - | - | 1 |
| *Ipomoea furcensis* Urb. | - | I | - | - | 1 |
| *Columnea sanguinea* Urb. | - | - | V | - | 1 |
| *Elaphoglossum crinitum* (L.) C. Chr. | - | - | V | - | 1 |
| *Elaphoglossum latifolium* (Sw.) J. Sm. | - | - | V | - | 1 |
| *Elleanthus cephalotus* Garay & Sweet | - | - | V | - | 1 |
| *Pleurothallis ruscifolia* (Jacq.) R. Br. | - | - | V | - | 1 |
| *Hyeronima domingensis* Urb. | - | - | V | - | 1 |
| *Calyptranthes selleanus* Urb. & Ekm. | - | - | V | - | 1 |
| *Torralbasia cuneifolia* (C. Wright) Krug. & Urb. | IV | - | III | - | 1 |
| *Meriania involucreta* (Desv.) Naud. | - | - | III | - | 1 |
| *Miconia punctata* (Desr.) D. Don | - | - | III | - | 1 |
| *Ophioglossum palmatum* L. | - | - | III | - | 1 |
| *Blechnum fragile* (Liebm.) Morton & Lellinger |  | - | III | - | 1 |
| *Arthrostylidium sarmentosum* Pilger | - | - | III | - | 1 |
| *Ilex tuerckheimii* Loes. | - | - | I | - | 1 |
| *Leandra limoides* (Urb.) W. Judd & Skee | - | - | I | - | 1 |
| *Maxillaria coccinea* (Jacq.) L.O. Wms. | - | - | I | - | 1 |
| *Asplenium radicans* L. | - | - | I | - | 1 |
| *Cestrum daphnoides* Griseb. | - | - | I | - | 1 |
| *Polypodium angustifolium* Sw. | - | - | I | - | 1 |
| *Ocotea foeniculacea* Mez | - | - | I | - | 1 |
| *Hillia parasitica* Jacq. | - | - | I | - | 1 |
| *Marattia kaulfussii* J. Smith | - | - | I | - | 1 |
| *Buchenavia tetraphylla* (Aubl.) R. A. Howard | - | - | - | V | 1 |
| *Byrsonima spicata* (Cav.) Kunth | - | - | - | V | 1 |
| *Casearea arborea* (L.C.Rich.) Urb. | - | - | - | V | 1 |
| *Clusia rosea* Jacq. | - | - | - | V | 1 |
| *Cyathea arborea* (L.) J.E. Smith | - | - | - | V | 1 |
| *Didymopanax morototoni* (Aubl.) Decne. & Planch | - | - | - | V | 1 |
| *Pytirogramma calomelanos* (L.) Link | - | - | - | V | 1 |
| *Miconia serrulata* (DC.) Naud. | - | - | - | V | 1 |
| *Ocotea globosa* (Aubl.) Schlecht. & Cham. | - | - | - | V | 1 |
| *Oreopanax capitatus* (Jacq.) Decne. & Planch. | - | - | - | V | 1 |
| *Ormosia krugii* Urb. | - | - | - | V | 1 |
| *Sloanea berteriana* Choisy | - | - | - | V | 1 |
| *Cnemidaria horrida* (L.) K. Presl | - | - | - | V | 1 |
| *Solanum torvum* Sw. | - | - | - | V | 1 |
| *Ipomoea tiliacea* (Willd.) Choisy | - | - | - | V | 1 |
| *Inga fagifolia* (L.) Willd. ex Benth. | - | - | - | III | 1 |
| *Inga vera* Willd. | - | - | - | III | 1 |
| *Bactris plumeriana* Mart. | - | - | - | III | 1 |
| *Nepsera aquatica* (Aubl.) Naud. | - | - | - | III | 1 |
| *Syngonium podophyllum* Schott | - | - | - | III | 1 |
| *Psychotria uliginosa* Sw. | - | - | - | III | 1 |
| *Urera baccifera* (L.) Gaud. | - | - | - | III | 1 |
| *Mimosa pudica* L. | - | - | - | I | 1 |
| *Neurolaena lobata* (L.) Cass. | - | - | - | I | 1 |
| *Triunfetta semitriloba* Jacq. | - | - | - | I | 1 |
| *Cupania americana* L. | - | - | - | I | 1 |
| *Hirtella triandra* Sw. | - | - | - | I | 1 |
| *Miconia racemosa* (Aubl.) DC. | - | - | - | I | 1 |
| *Zantoxylum martinicensis* (Lam.) DC. | - | - | - | I | 1 |
| *Cissus verticillata* (L.) Nicholson & Farris | - | - | - | I | 1 |
| *Entada gigas* (L.) Fawc. & Rendle | - | - | - | I | 1 |
| *Palicourea crocea* (Sw.) Schultes | - | - | - | I | 1 |
| *Piper adunculum* L. | - | - | - | I | 1 |
| *Coccocypselum herbaceum* Aubl. | - | - | - | I | 1 |
| *Hedychium coronarium* Koen. | - | - | - | I | 1 |
| *Solanum jamaicense* Mill. | - | - | - | I | 1 |
| *Urena lobata* L. | - | - | - | I | 1 |

As1. *Hyeronimo montanae*-*Magnolietum pallescenti*. As2. *Cyathea furfuracei*-*Prestoetum montanae*. As3. *Hyeronimo dominguensis*-*Magnolietum hamorii*. As4. *Ormosia krugii*-*Prestoetum montanae*. P. Presences.

### Conservation status of the associations

The analysis of the floristic diversity of the relevés shows a predominance of Shannon_T diversity (total diversity) over the diversity of non-endemic and endemic species, except in the samplings DR15, DR16 and DR17, where there is a coincidence between Shannon_T and Shannon_Ne due to the low rate of endemic species, with only two species: *Bactris plumeriana* and *Clidemia umbellata*. The diversity rate for characteristic species (Shannon_Ca) tends to be high compared to companion species (Shannon_Co), except in DR3 which has a value for Shannon_Co = 1.099 (Table 5).

**Table 5.** Shannon diversity by relevé.

|  | DR1 | DR2 | DR3 | DR4 | DR5 | DR6 | DR7 | DR8 | DR9 | DR10 | DR11 | DR12 | DR13 | DR14 | DR15 | DR16 | DR17 |
| --- | --- | --- | --- | --- | --- | --- | --- | --- | --- | --- | --- | --- | --- | --- | --- | --- | --- |
| Shannon_T | 3,612 | 3,443 | 2,451 | 3,566 | 3,464 | 3,557 | 3,557 | 3,458 | 3,424 | 3,389 | 3,786 | 3,972 | 3,781 | 3,702 | 3,917 | 3,496 | 3,499 |
| Shannon_Ca | 3,17 | 3,247 | 2,165 | 3,097 | 3,119 | 3,152 | 2,819 | 2,606 | 2,803 | 2,563 | 3,119 | 3,154 | 3,013 | 2,683 | 3,173 | 3,061 | 3,027 |
| Shannon_Co | 2,591 | 1,718 | 1,099 | 2,602 | 2,232 | 2,507 | 2,91 | 2,901 | 2,655 | 2,814 | 3,066 | 3,391 | 3,162 | 3,256 | 3,277 | 2,458 | 2,532 |
| Shannon_Ne | 3,178 | 2,947 | 2,16 | 3 | 2,997 | 3,119 | 3,347 | 3,244 | 3,104 | 2,998 | 3,561 | 3,601 | 3,388 | 3,458 | 3,897 | 3,435 | 3,438 |
| Shannon_E | 2,574 | 2,508 | 1,089 | 2,732 | 2,49 | 2,557 | 1,891 | 1,842 | 2,137 | 2,158 | 2,41 | 2,803 | 2,658 | 2,074 | 0 | 0 | 0 |

(Shannon\_T= total diversity; Shannon\_Ca= characteristic community species diversity; Shannon\_Co= companion community species diversity; Shannon\_Ne= non-endemic species diversity; Shannon\_E= endemic species diversity).

In the comparative analysis of the diversity between the four associations using the average diversity values for each relevé, it can be seen that association As4 has a Shannon_E =0 due to an almost total lack of endemic species. This association also has low values for total diversity and non-endemic species, with 44.2% trees, 22.9% shrubs, 13.1% climbing plants and 16.3% herbs; whereas the other associations have a greater diversity. The Shannon_Ca value is higher than Shannon_Co in the four associations except for As3; however, the values are similar due to a tendency to ingression by companion species from neighbouring communities (Table 6, Fig 6).

**Fig 6.**
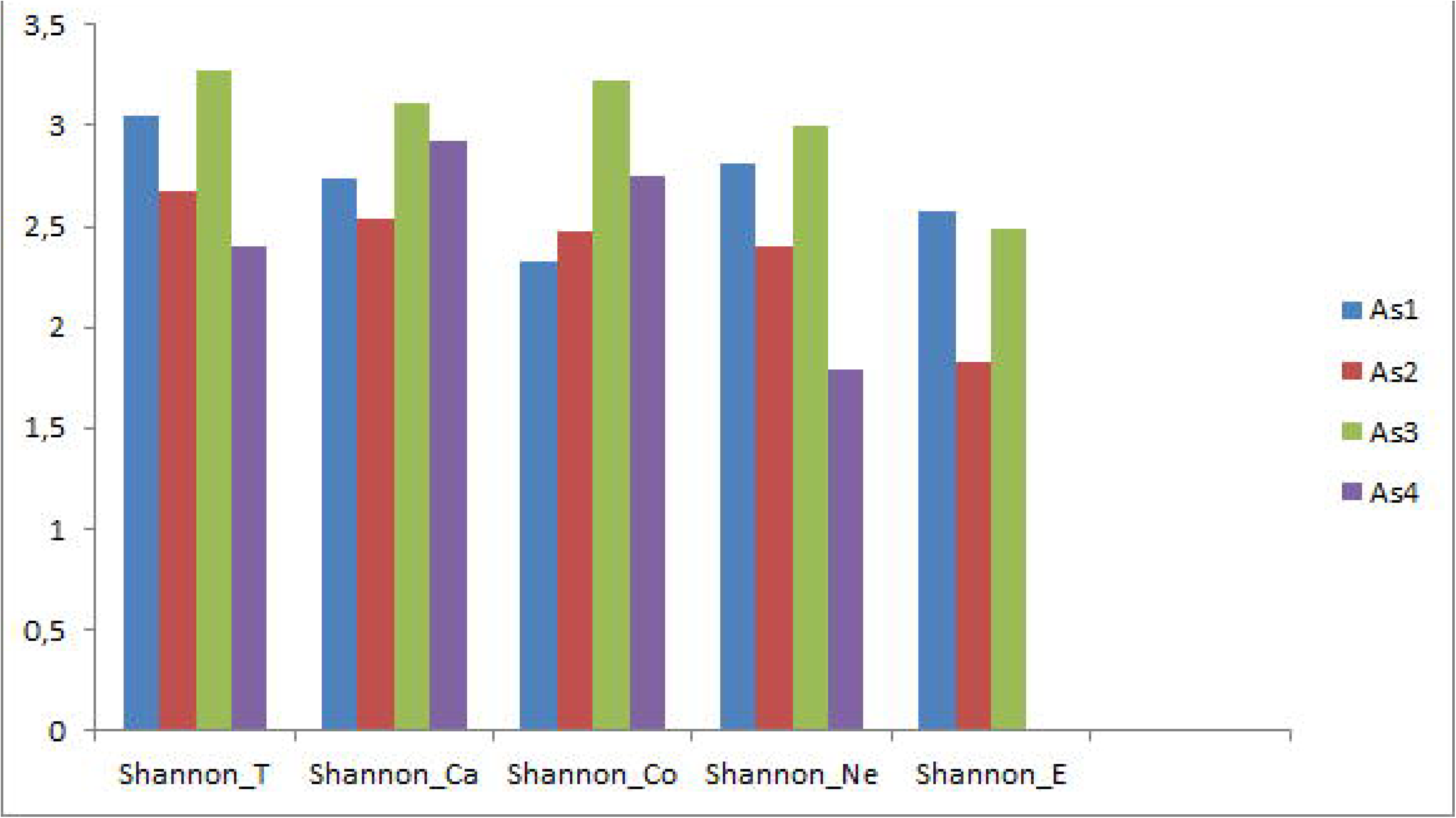
Shannon diversity value (T, Ca, Co, Ne, E). As1. *Hyeronimo montanae-Magnolietum pallescentis*. As2. *Cyatheo furfuracei-Prestoetum motanae*. As3. *Hyeronimo dominguensis-Magnolietum hamorii.* As4. *Ormosio krugii-Prestoetum montanae*.

**Table 6.**
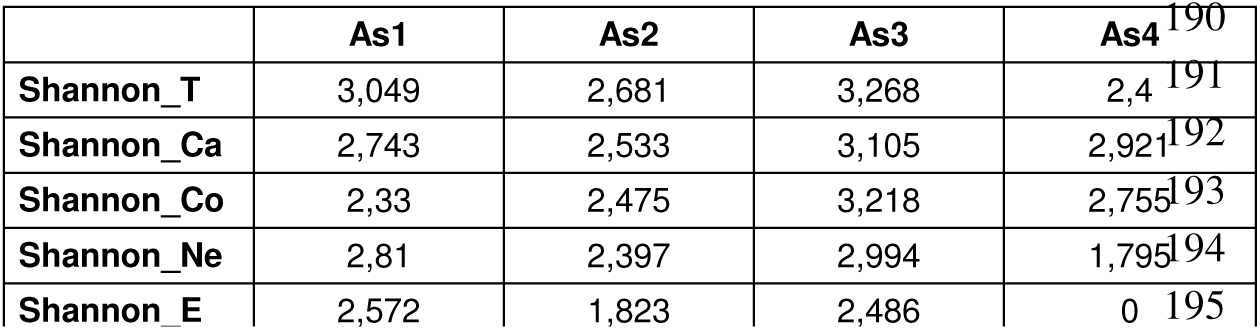
Diversity analysis of each of the four plant associations.

## Discussion

In all cases there is a high diversity of trees, among which it is particularly worth noting the endemics *Magnolia pallescens* Urb. & Ekm., *Hyeronima montana* A. Liogier, *Magnolia hamorii* Howard, *Hyeronima domingensis* Urb., *Malpighia macracantha* Ekm. & Nied., and *Bactris plumeriana* Mart. These are therefore plant communities with an endemic character that require protection measures. Although all four associations are of great interest to conservation, the two best conserved associations have the highest rate of endemics, and are precisely the ones located in the Bahoruco-Hottense and central biogeographical sectors [18,20] which concurs with the floristic studies of [1,3,11,43]. However, the areas exposed to greater environmental impact, as is the case of biogeographical sectors such as the Cordillera Oriental range which are subjected to significant human pressure, have less floristic diversity and a lower number of endemic species. No significant differences can be seen between the relevés in the Shannon diversity index, whose values range between DR3 with indexes of Sh = 2.451, and DR12 with higher values of Sh = 3.972 (Table 5); this does not imply that DR3 is poorly conserved [27], but simply that there is an almost complete predominance of the faithful species *Prestoea montana*, which has a high cover and very few companion species. However, relevé DR12 contains many individuals with low cover and a high rate of companion species. The low rate of endemisms in As4 represented by relevés DR15, DR16, DR17 in the Cordillera Oriental range is the result of significant anthropic action owing to population density.

The four associations described are included in the phytosociological classes *Weinmannio-Cyrilletea* Knapp 1964 and *Ocoteo-Magnolietea* Borhidi and Muñiz, in Borhidi et al. 1979. Due to the high floristic and biogeographical differentiation between Hispaniola and Cuba (Tables 2 and 3), these associations cannot be included in any of the alliances described for the island of Cuba. We therefore propose two new alliances: all. *Rondeletio ochraceae-Clusion roseae*, in which the alliance species are *Rondeletia ochracea, Turpinia occidentalis, Clusia rosea, Mikania cordifolia, Alchornea latifolia*, and *Cyatheo furfuracei-Prestoetum motanae* is the type association; and all. *Rondeletio ochraceae-Didymopanion tremuli*, with the species *Rondeletia ochracea, Didymopanax tremulus, Psychotria guadalupensis,* and *Hyeronimo montanae-Magnolietum pallescentis* as the type association. All these results are according to [44]

## Conclusions

This study in the Dominican Republic reveals the existence of different types of rainforest that are clearly differentiated by their floristic, biogeographical and bioclimatic composition. This broadleaved forest or rainforest is frequent in the Sierra de Bahoruco and the Cordillera Central, Septentrional and Oriental ranges due to the increased rainfall in these areas caused by the impact of moisture-laden Atlantic winds. Differences in soil and biogeography have conditioned a rich and different flora. The Cordillera Central range –geologically the oldest, and with a siliceous character– is home to rainforests of *Magnolia pallescens* and forests of *Prestoea montana* (As1 and As2) in humid-hyper-humid areas; whereas the associations As3 in Bahoruco and As4 in the Cordillera Oriental range also develop in humid environments but on soil substrates. This leads us to propose four new syntaxa with the rank of association and two new alliances.

## Syntaxonomical checklist for the cloud forest of Hispaniola

*Weinmannio-Cyrilletea* Knapp 1964

*Weinmannio-Cyrilletalia* Knapp 1964

*Rondeletio ochraceae-Clusion roseae* Cano, Cano-Ortiz & Veloz all. nova hoc loco

*Cyatheo furfuracei-Prestoetum motanae* Cano, Cano-Ortiz & Veloz ass. nova hoc loco

*Ormosio krugii-Prestoetum montanae* Cano, Cano-Ortiz & Veloz ass. nova hoc loco

*Ocoteo-Magnolietea* Borhidi and Muñiz in Borhdi et al. 1979

*Ocoteo-Magnolietalia* Muñiz in Borhdi et al. 1979

*Rondeletio ochraceae-Didymopanion tremuli* Cano, Cano-Ortiz & Veloz all. nova hoc loco

*Hyeronimo montanae-Magnolietum pallescentis* Cano, Cano-Ortiz & Veloz ass. nova hoc loco

*Hyeronimo dominguensis-Magnolietum hamorii* Cano, Cano-Ortiz & Veloz ass. nova hoc loco

## Acknowledgments

We are very grateful to Ms Pru Brooke Turner (MA Cantab.) for the English translation of this article. This manuscript has been released as a Pre-Print at: Biorxiv 543892; doi: https://doi.org/10.1101/5438922019.

---

**S1 Table 1.**
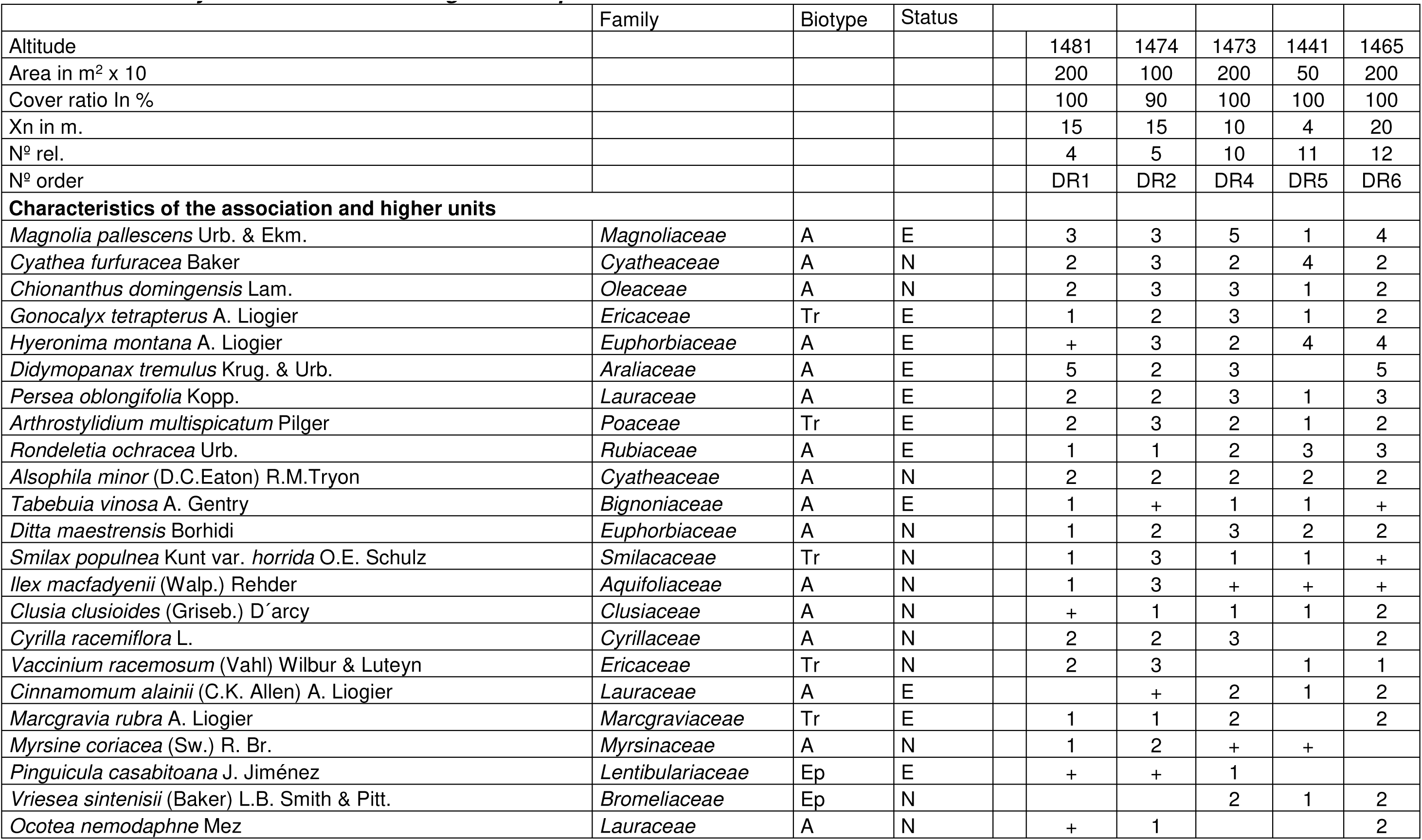
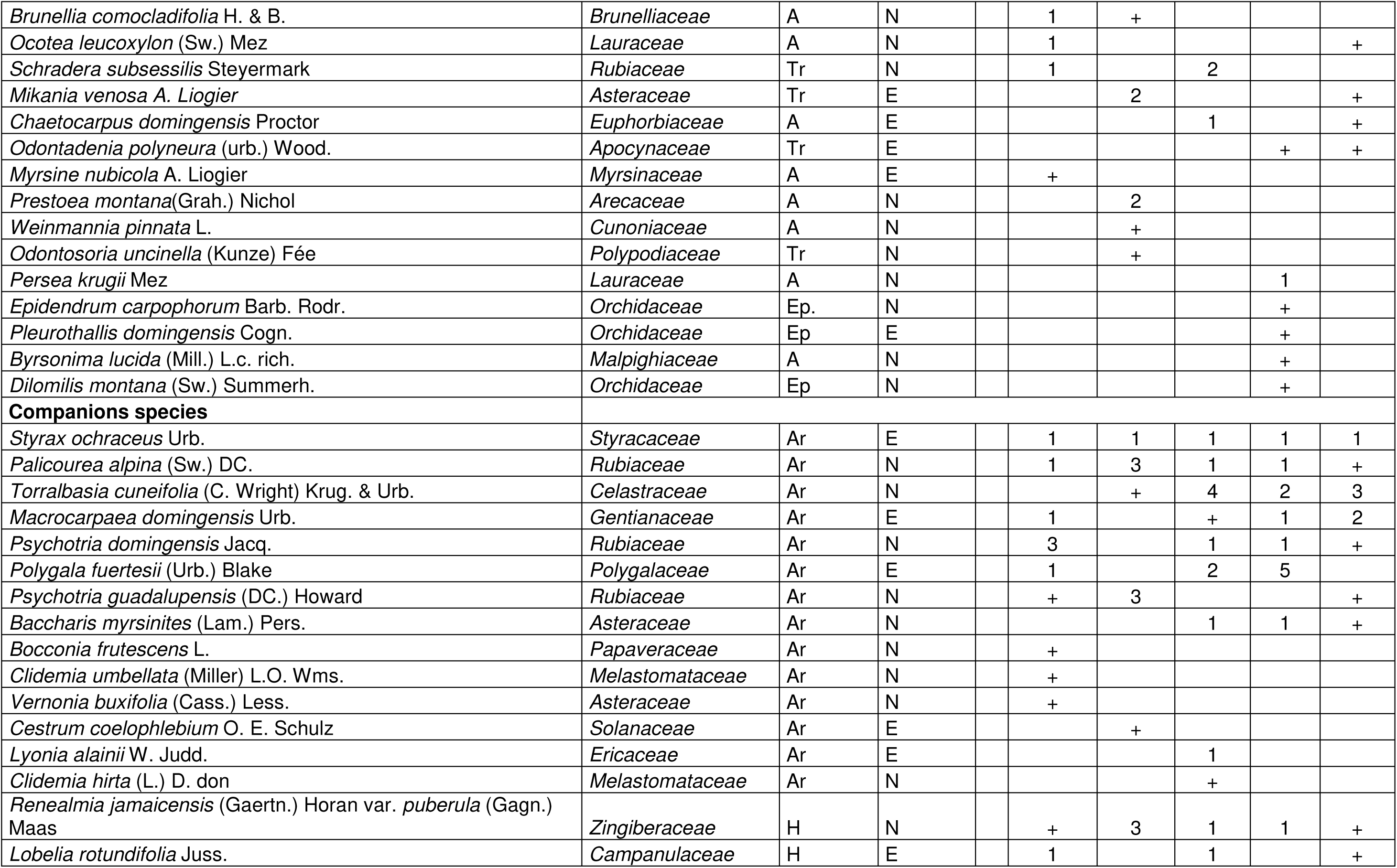
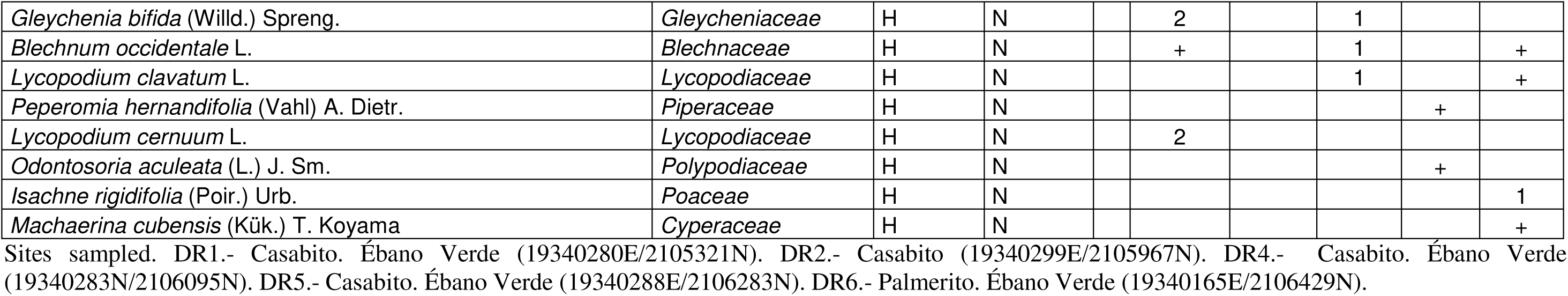
Ass. Hyeronimo montanae-Magnolietum pallescentis.

**S2 Table 2.**
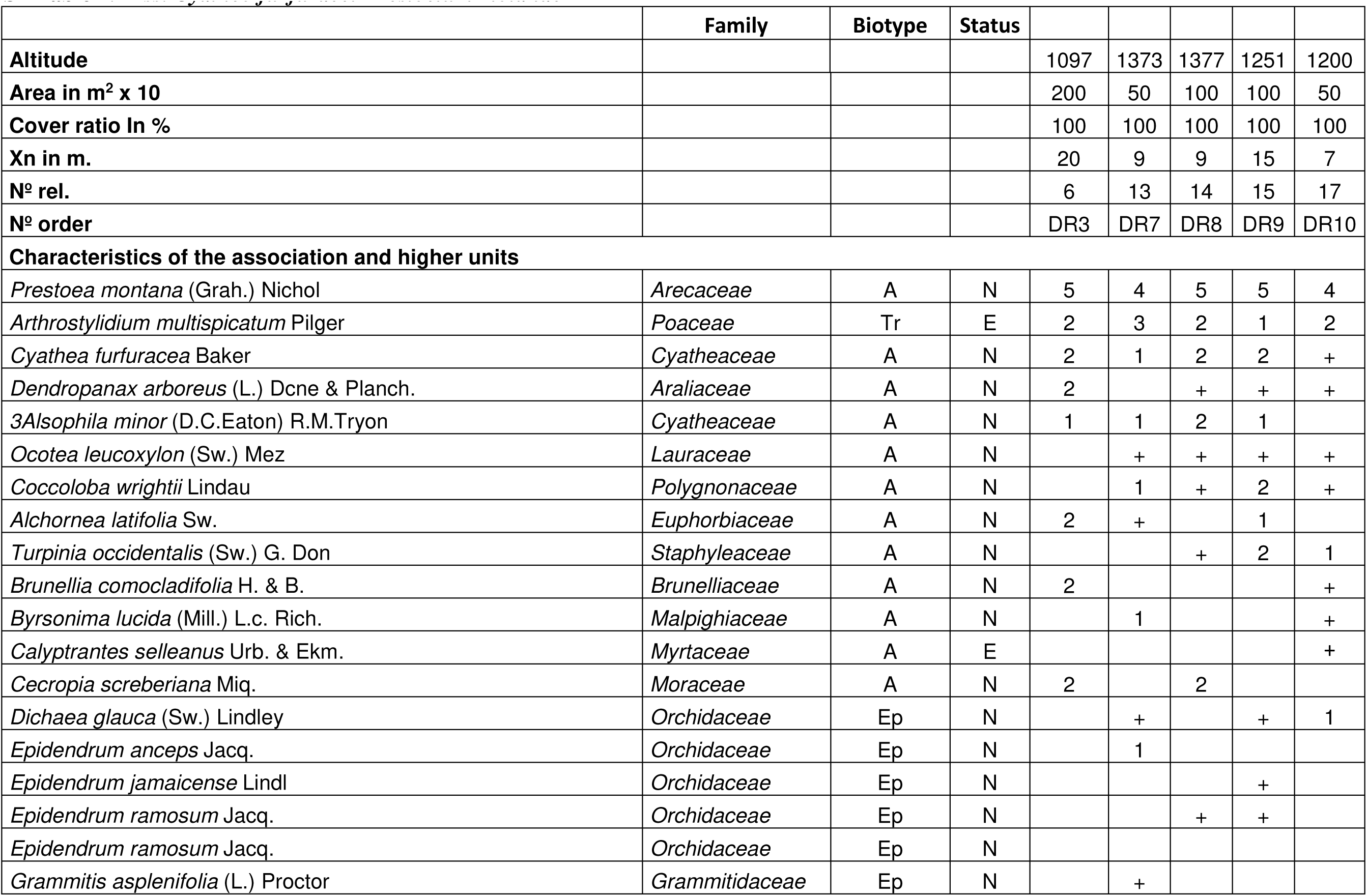
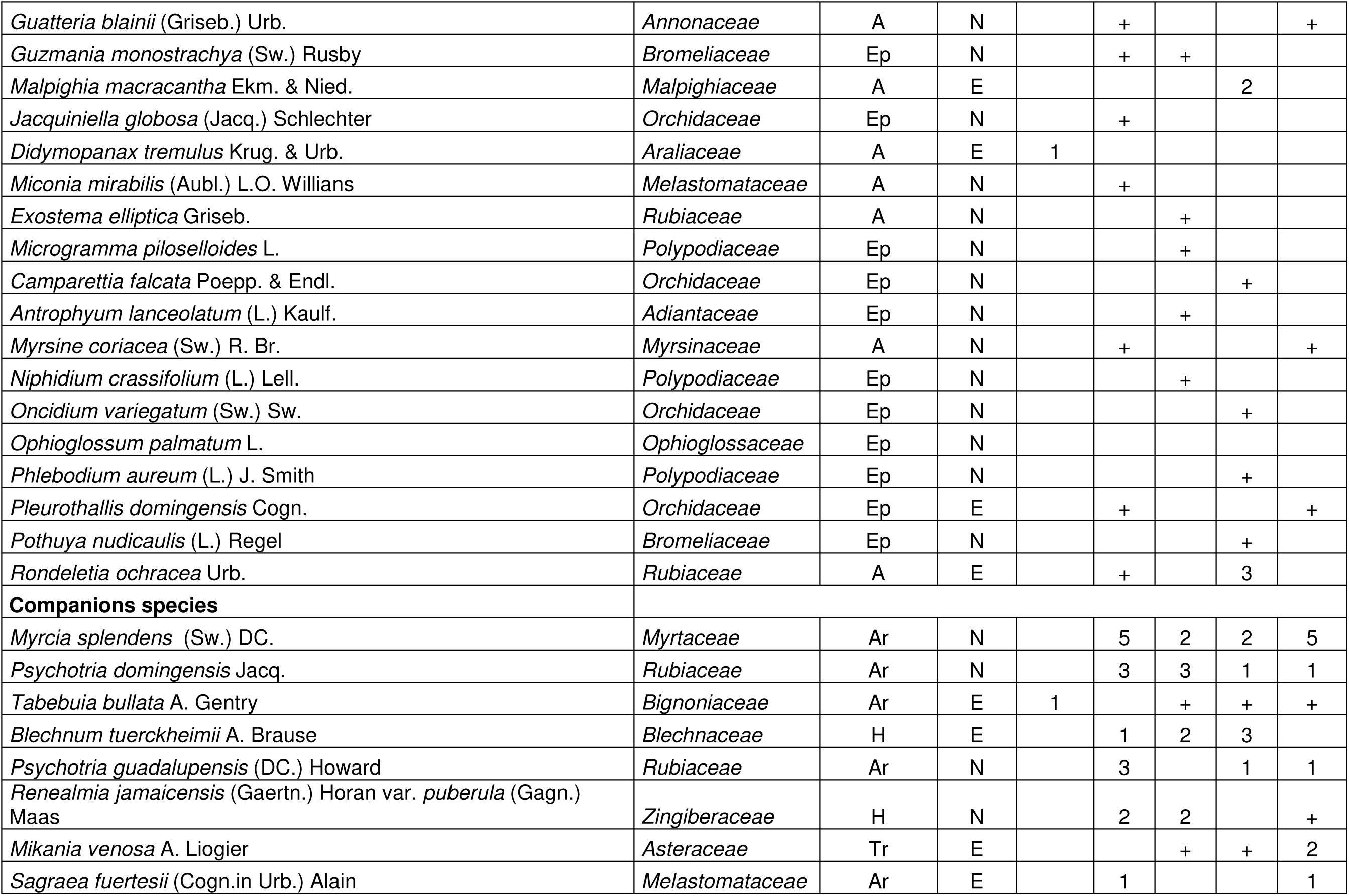
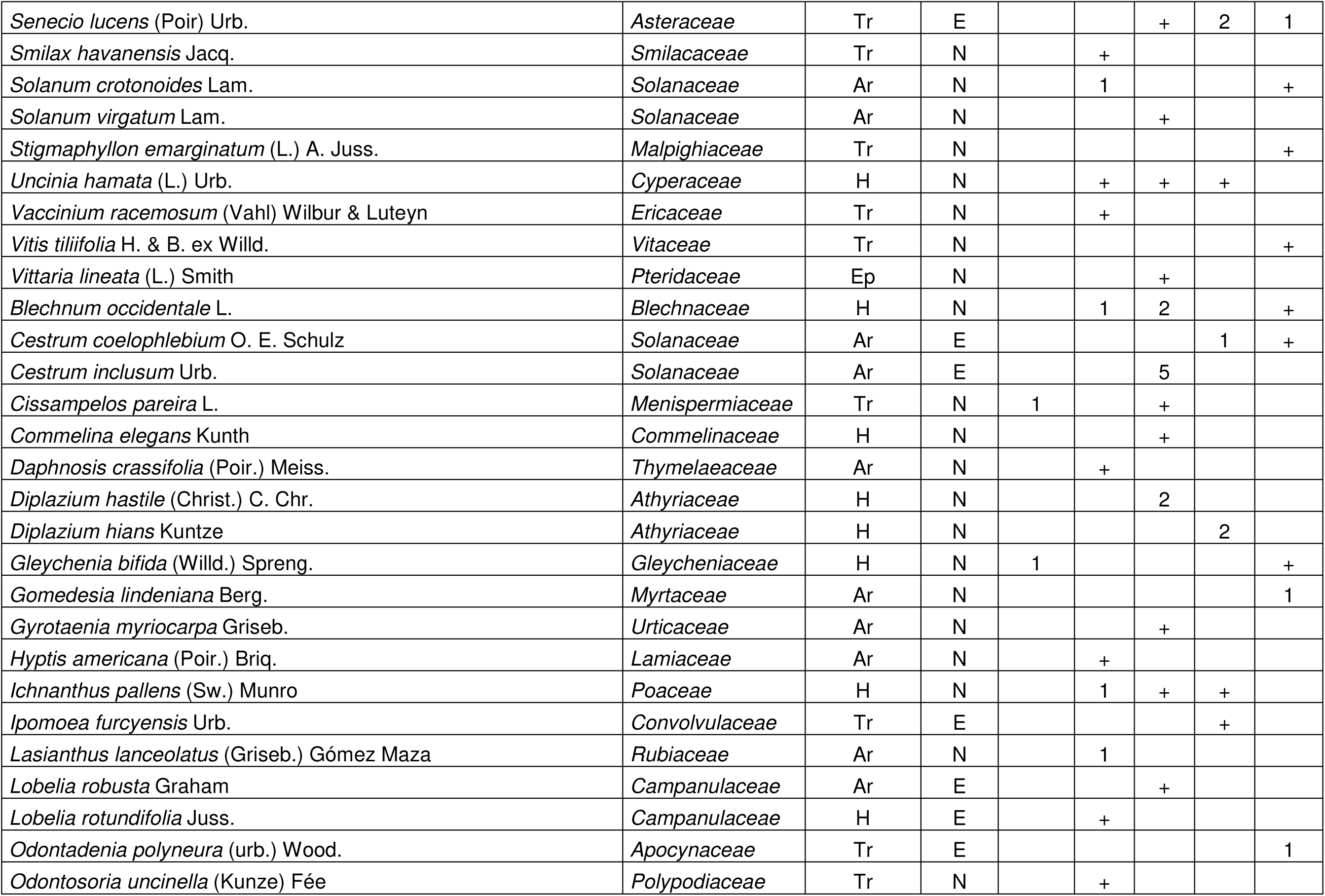
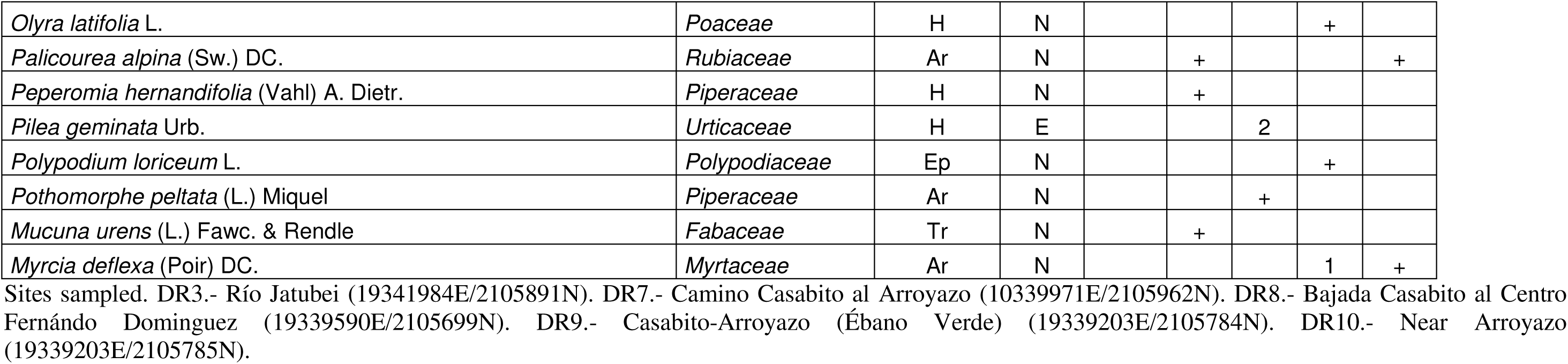
Ass. Cyatheo furfuracei-Prestoetum motanae.

**S3 Table 3.**
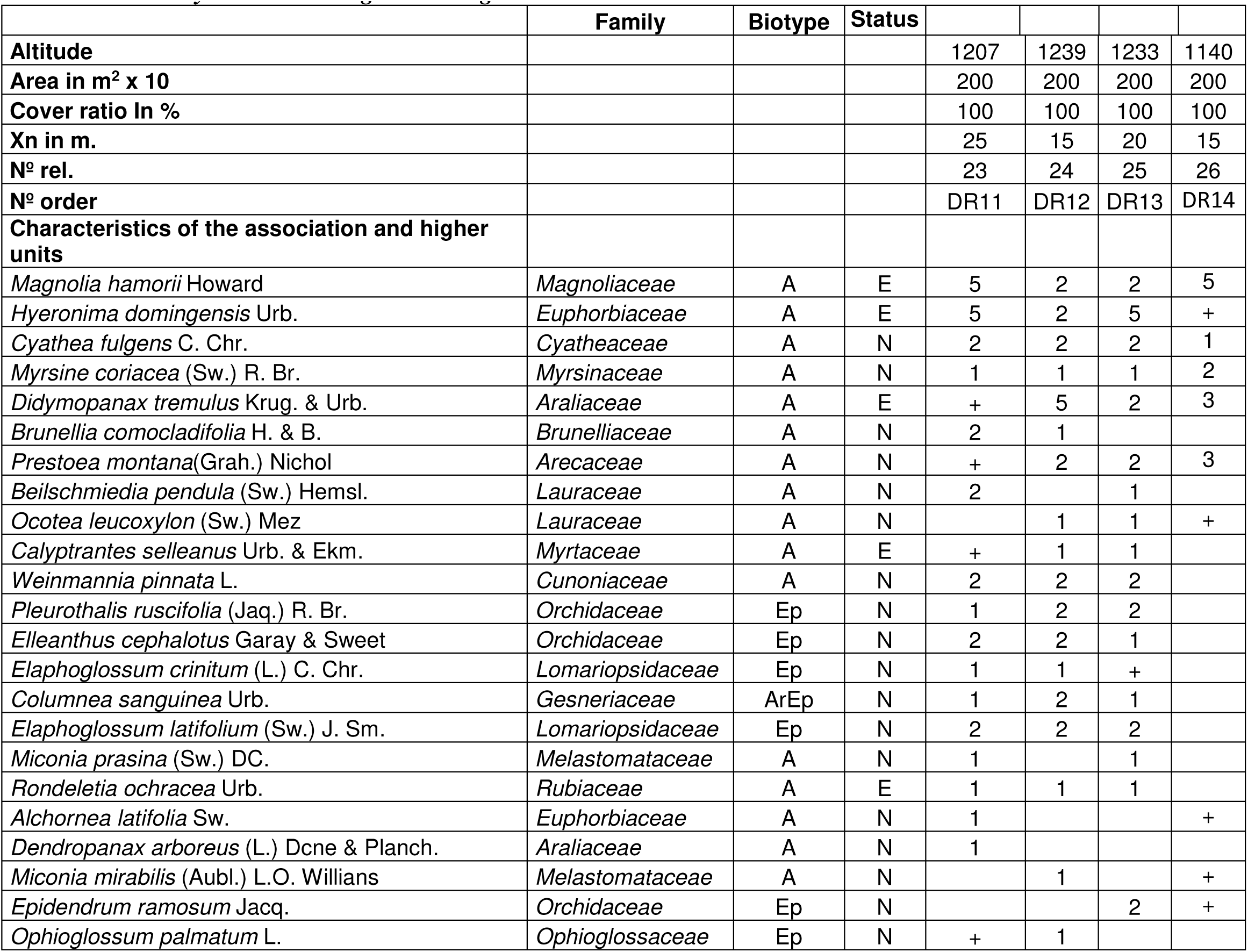
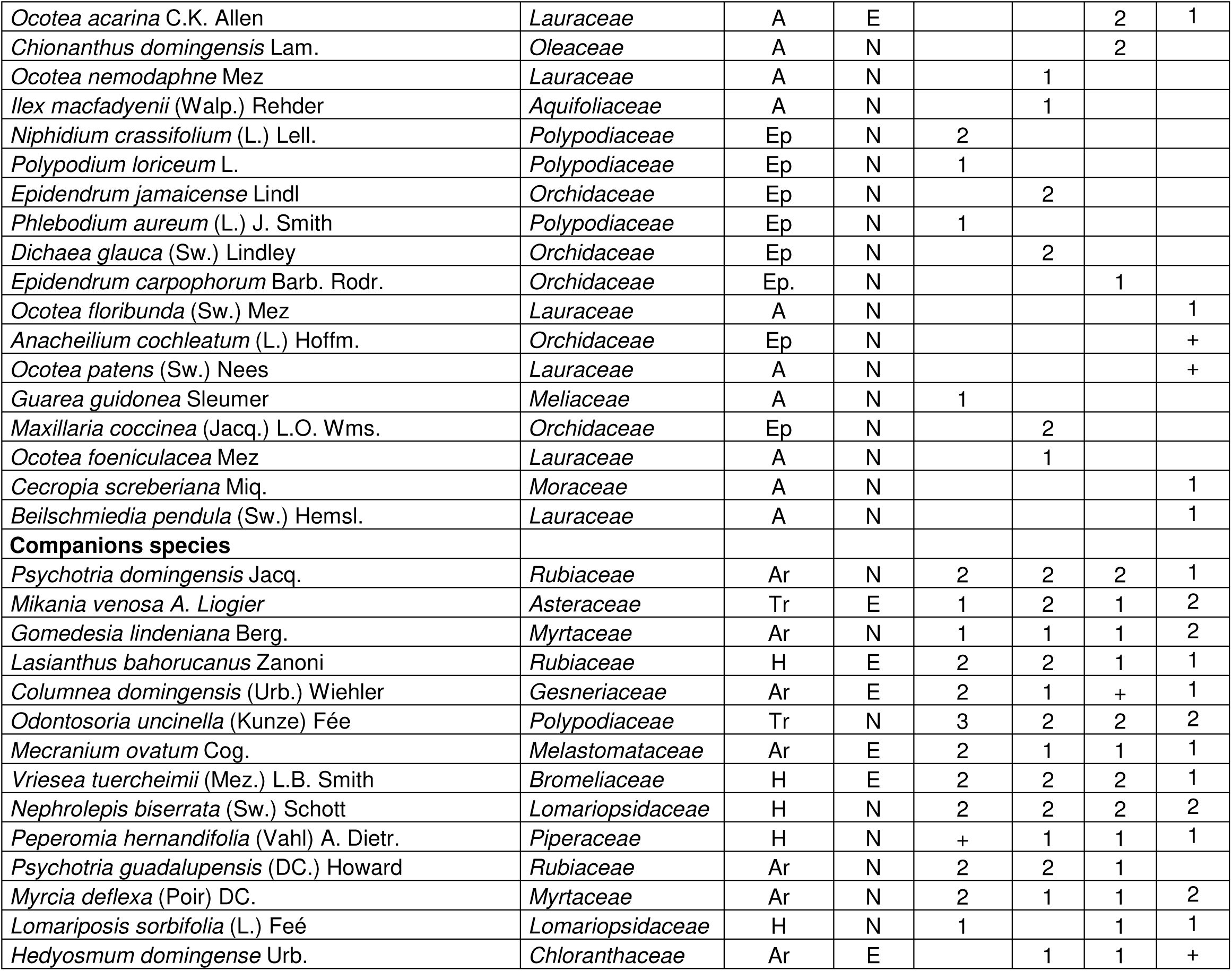
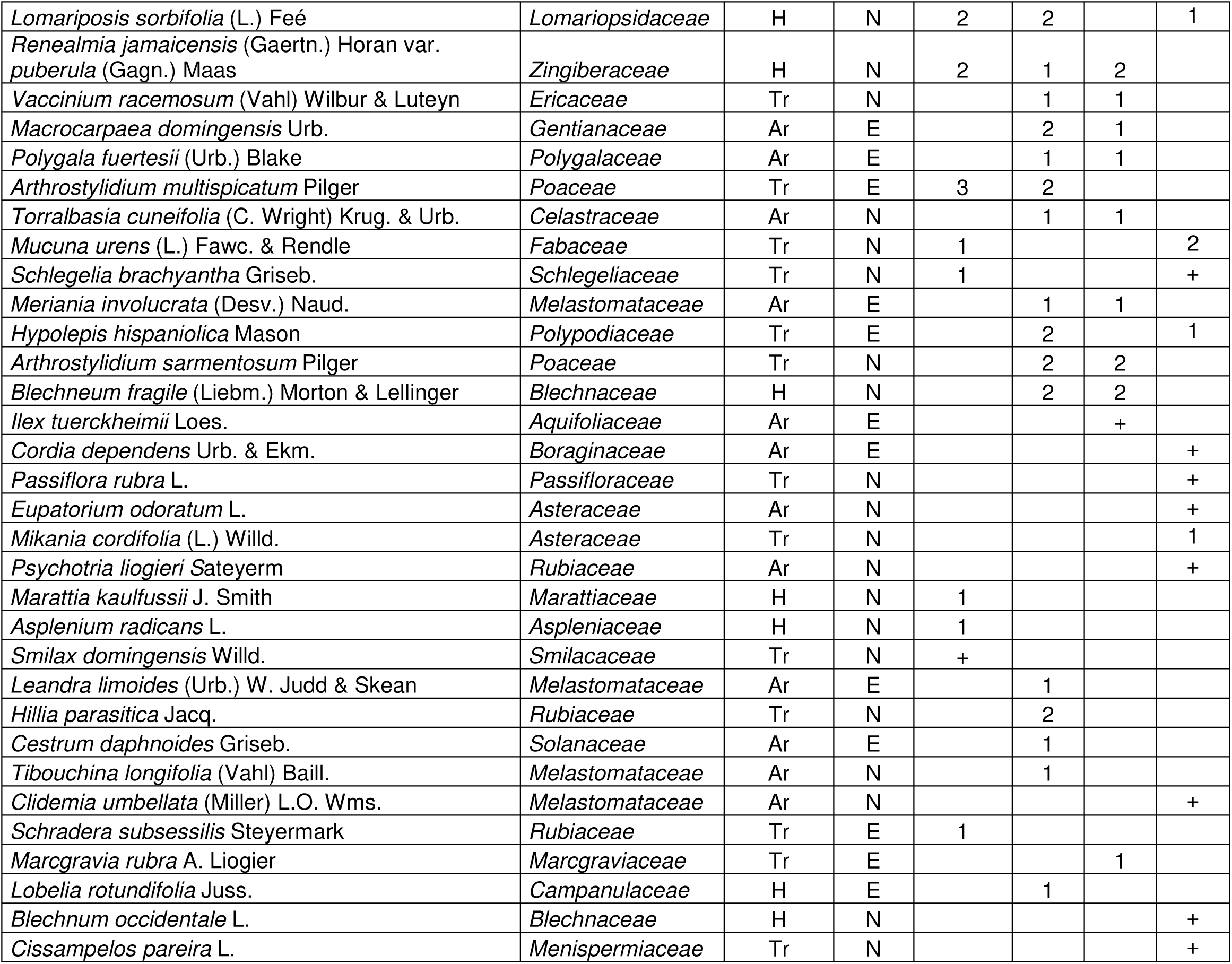
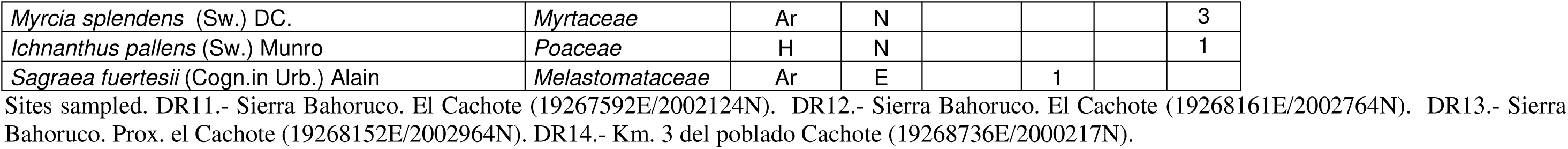
Ass. Hyeronimo dominguensis-Magnolietum hamorii.

**S4 Table 4.**
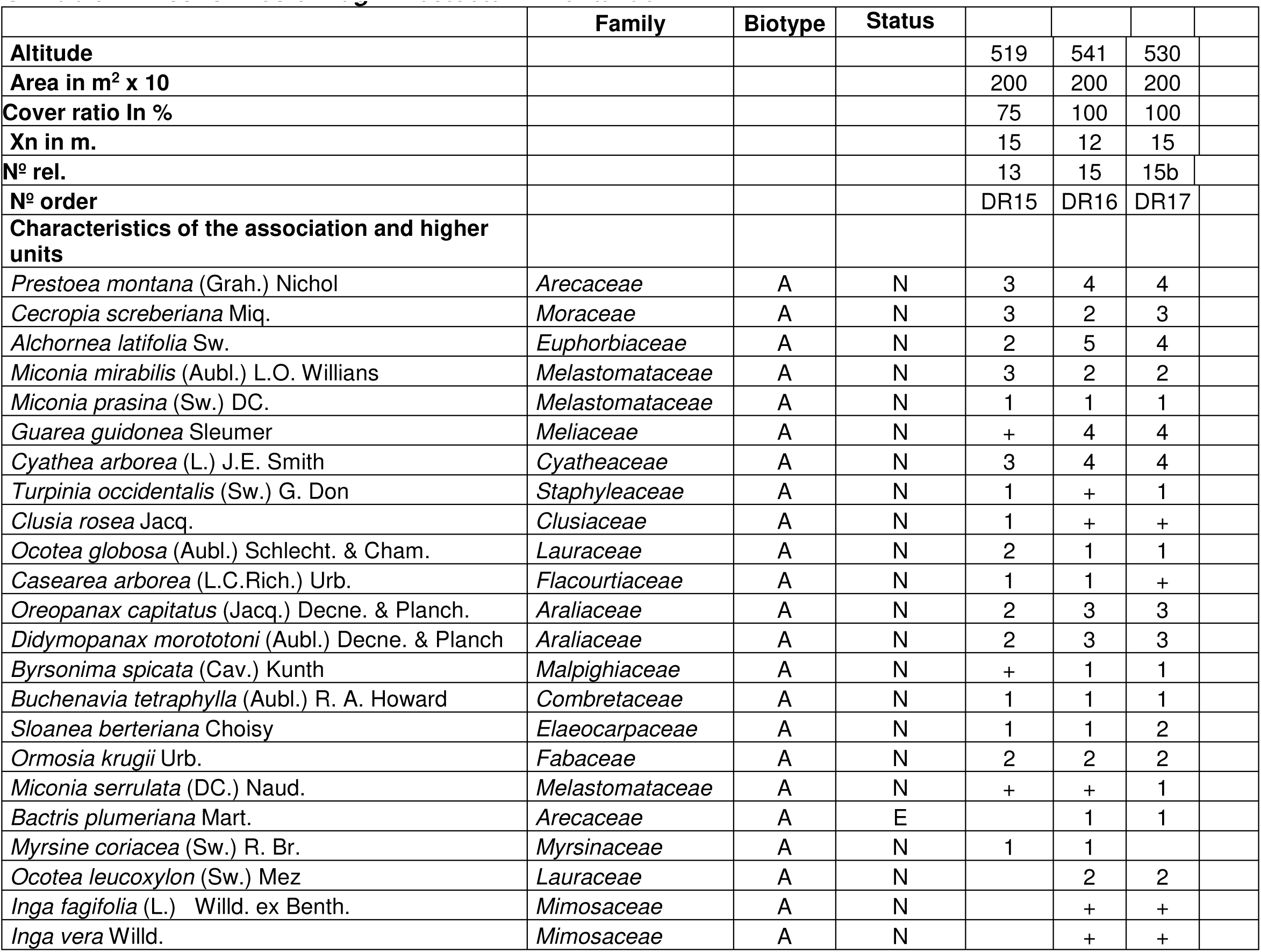
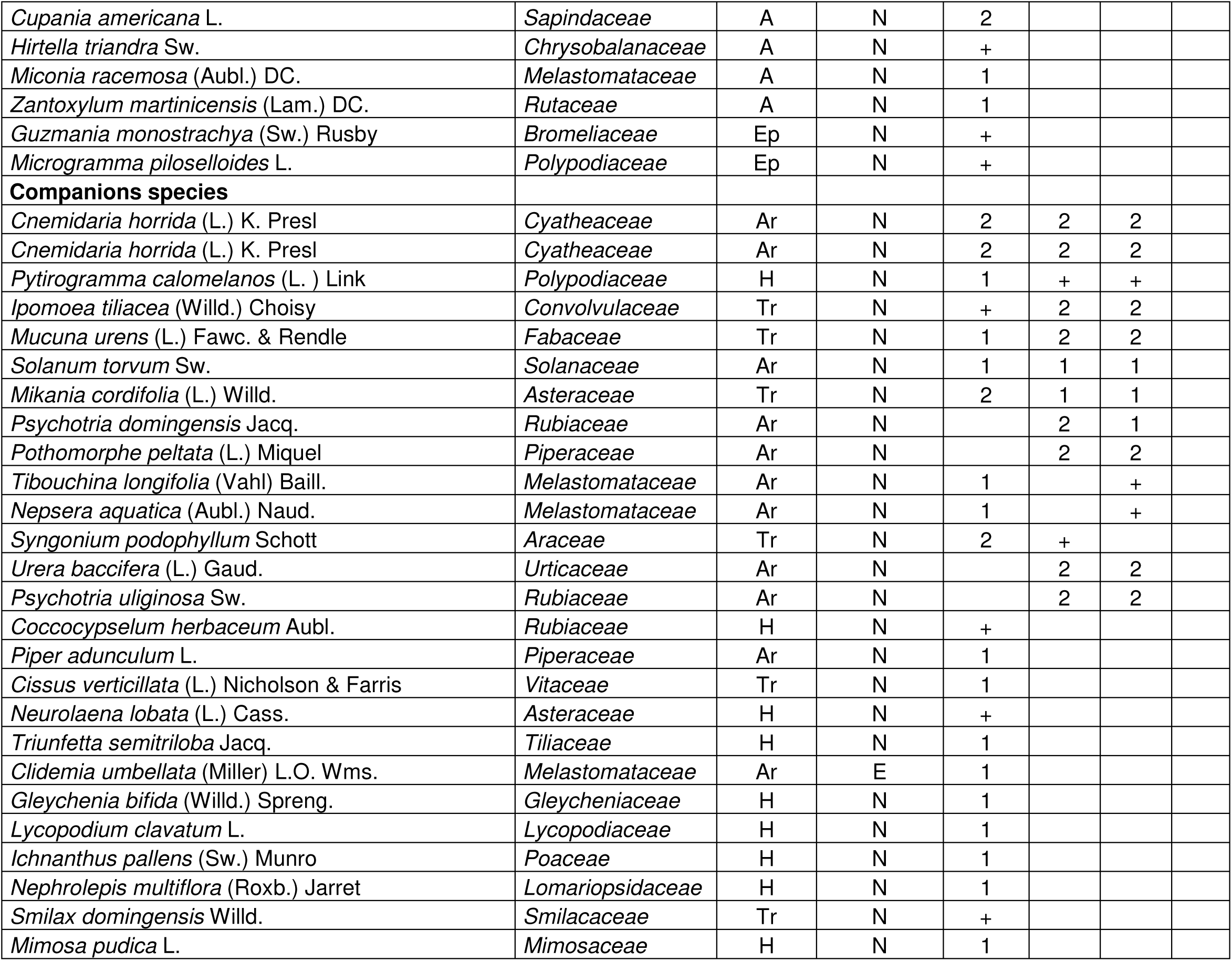
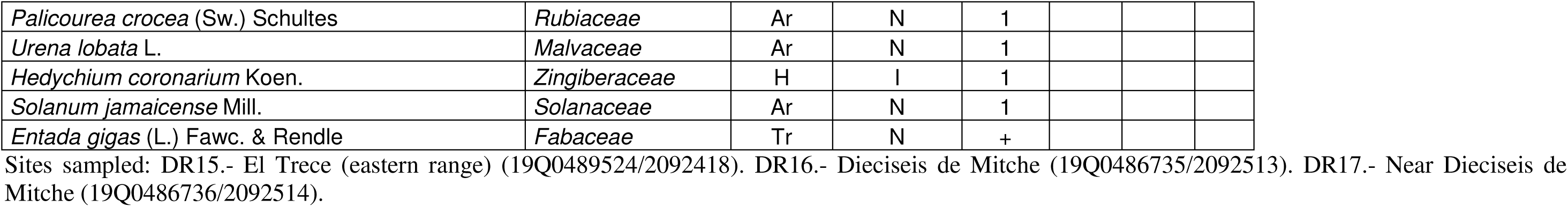
Ass. Ormosio krugii-Prestoetum montanae.

