## Supplemental file 1 for "The cloud forest in the Dominican Republic: diversity and conservation status"

**S1 Table 1. *Ass. Hyeronimo montanae-Magnolietum pallescentis*.**

|  | Family | Biotype | Status |  | |  |  |  |  |
| --- | --- | --- | --- | --- | --- | --- | --- | --- | --- |
| Altitude |  |  |  |  | 1481 | 1474 | 1473 | 1441 | 1465 |
| Area in m^2^ x 10 |  |  |  |  | 200 | 100 | 200 | 50 | 200 |
| Cover ratio In % |  |  |  |  | 100 | 90 | 100 | 100 | 100 |
| Xn in m. |  |  |  |  | 15 | 15 | 10 | 4 | 20 |
| Nº rel. |  |  |  |  | 4 | 5 | 10 | 11 | 12 |
| Nº order |  |  |  |  | DR1 | DR2 | DR4 | DR5 | DR6 |
| **Characteristics of the association and higher units** | |  |  |  |  |  |  |  |  |
| *Magnolia pallescens* Urb. & Ekm. | *Magnoliaceae* | A | E |  | 3 | 3 | 5 | 1 | 4 |
| *Cyathea furfuracea* Baker | *Cyatheaceae* | A | N |  | 2 | 3 | 2 | 4 | 2 |
| *Chionanthus domingensis* Lam. | *Oleaceae* | A | N |  | 2 | 3 | 3 | 1 | 2 |
| *Gonocalyx tetrapterus* A. Liogier | *Ericaceae* | Tr | E |  | 1 | 2 | 3 | 1 | 2 |
| *Hyeronima montana* A. Liogier | *Euphorbiaceae* | A | E |  | + | 3 | 2 | 4 | 4 |
| *Didymopanax tremulus* Krug. & Urb. | *Araliaceae* | A | E |  | 5 | 2 | 3 |  | 5 |
| *Persea oblongifolia* Kopp. | *Lauraceae* | A | E |  | 2 | 2 | 3 | 1 | 3 |
| *Arthrostylidium multispicatum* Pilger | *Poaceae* | Tr | E |  | 2 | 3 | 2 | 1 | 2 |
| *Rondeletia ochracea* Urb. | *Rubiaceae* | A | E |  | 1 | 1 | 2 | 3 | 3 |
| *Alsophila minor* (D.C.Eaton) R.M.Tryon | *Cyatheaceae* | A | N |  | 2 | 2 | 2 | 2 | 2 |
| *Tabebuia vinosa* A. Gentry | *Bignoniaceae* | A | E |  | 1 | + | 1 | 1 | + |
| *Ditta maestrensis* Borhidi | *Euphorbiaceae* | A | N |  | 1 | 2 | 3 | 2 | 2 |
| *Smilax populnea* Kunt var. *horrida* O.E. Schulz | *Smilacaceae* | Tr | N |  | 1 | 3 | 1 | 1 | + |
| *Ilex macfadyenii* (Walp.) Rehder | *Aquifoliaceae* | A | N |  | 1 | 3 | + | + | + |
| *Clusia clusioides* (Griseb.) D´arcy | *Clusiaceae* | A | N |  | + | 1 | 1 | 1 | 2 |
| *Cyrilla racemiflora* L. | *Cyrillaceae* | A | N |  | 2 | 2 | 3 |  | 2 |
| *Vaccinium racemosum* (Vahl) Wilbur & Luteyn | *Ericaceae* | Tr | N |  | 2 | 3 |  | 1 | 1 |
| *Cinnamomum alainii* (C.K. Allen) A. Liogier | *Lauraceae* | A | E |  |  | + | 2 | 1 | 2 |
| *Marcgravia rubra* A. Liogier | *Marcgraviaceae* | Tr | E |  | 1 | 1 | 2 |  | 2 |
| *Myrsine coriacea* (Sw.) R. Br. | *Myrsinaceae* | A | N |  | 1 | 2 | + | + |  |
| *Pinguicula casabitoana* J. Jiménez | *Lentibulariaceae* | Ep | E |  | + | + | 1 |  |  |
| *Vriesea sintenisii* (Baker) L.B. Smith & Pitt. | *Bromeliaceae* | Ep | N |  |  |  | 2 | 1 | 2 |
| *Ocotea nemodaphne* Mez | *Lauraceae* | A | N |  | + | 1 |  |  | 2 |
| *Brunellia comocladifolia* H. & B. | *Brunelliaceae* | A | N |  | 1 | + |  |  |  |
| *Ocotea leucoxylon* (Sw.) Mez | *Lauraceae* | A | N |  | 1 |  |  |  | + |
| *Schradera subsessilis* Steyermark | *Rubiaceae* | Tr | N |  | 1 |  | 2 |  |  |
| *Mikania venosa A. Liogier* | *Asteraceae* | Tr | E |  |  | 2 |  |  | + |
| *Chaetocarpus domingensis* Proctor | *Euphorbiaceae* | A | E |  |  |  | 1 |  | + |
| *Odontadenia polyneura* (urb.) Wood. | *Apocynaceae* | Tr | E |  |  |  |  | + | + |
| *Myrsine nubicola* A. Liogier | *Myrsinaceae* | A | E |  | + |  |  |  |  |
| *Prestoea montana*(Grah.) Nichol | *Arecaceae* | A | N |  |  | 2 |  |  |  |
| *Weinmannia pinnata* L. | *Cunoniaceae* | A | N |  |  | + |  |  |  |
| *Odontosoria uncinella* (Kunze) Fée | *Polypodiaceae* | Tr | N |  |  | + |  |  |  |
| *Persea krugii* Mez | *Lauraceae* | A | N |  |  |  |  | 1 |  |
| *Epidendrum carpophorum* Barb. Rodr. | *Orchidaceae* | Ep. | N |  |  |  |  | + |  |
| *Pleurothallis domingensis* Cogn. | *Orchidaceae* | Ep | E |  |  |  |  | + |  |
| *Byrsonima lucida* (Mill.) L.c. rich. | *Malpighiaceae* | A | N |  |  |  |  | + |  |
| *Dilomilis montana* (Sw.) Summerh. | *Orchidaceae* | Ep | N |  |  |  |  | + |  |
| **Companions species** |  | | | | | | | | |
| *Styrax ochraceus* Urb. | *Styracaceae* | Ar | E |  | 1 | 1 | 1 | 1 | 1 |
| *Palicourea alpina* (Sw.) DC. | *Rubiaceae* | Ar | N |  | 1 | 3 | 1 | 1 | + |
| *Torralbasia cuneifolia* (C. Wright) Krug. & Urb. | *Celastraceae* | Ar | N |  |  | + | 4 | 2 | 3 |
| *Macrocarpaea domingensis* Urb. | *Gentianaceae* | Ar | E |  | 1 |  | + | 1 | 2 |
| *Psychotria domingensis* Jacq. | *Rubiaceae* | Ar | N |  | 3 |  | 1 | 1 | + |
| *Polygala fuertesii* (Urb.) Blake | *Polygalaceae* | Ar | E |  | 1 |  | 2 | 5 |  |
| *Psychotria guadalupensis* (DC.) Howard | *Rubiaceae* | Ar | N |  | + | 3 |  |  | + |
| *Baccharis myrsinites* (Lam.) Pers. | *Asteraceae* | Ar | N |  |  |  | 1 | 1 | + |
| *Bocconia frutescens* L. | *Papaveraceae* | Ar | N |  | + |  |  |  |  |
| *Clidemia umbellata* (Miller) L.O. Wms. | *Melastomataceae* | Ar | N |  | + |  |  |  |  |
| *Vernonia buxifolia* (Cass.) Less. | *Asteraceae* | Ar | N |  | + |  |  |  |  |
| *Cestrum coelophlebium* O. E. Schulz | *Solanaceae* | Ar | E |  |  | + |  |  |  |
| *Lyonia alainii* W. Judd. | *Ericaceae* | Ar | E |  |  |  | 1 |  |  |
| *Clidemia hirta* (L.) D. don | *Melastomataceae* | Ar | N |  |  |  | + |  |  |
| *Renealmia jamaicensis* (Gaertn.) Horan var. *puberula* (Gagn.) Maas | *Zingiberaceae* | H | N |  | + | 3 | 1 | 1 | + |
| *Lobelia rotundifolia* Juss. | *Campanulaceae* | H | E |  | 1 |  | 1 |  | + |
| *Gleychenia bifida* (Willd.) Spreng. | *Gleycheniaceae* | H | N |  | 2 |  | 1 |  |  |
| *Blechnum occidentale* L. | *Blechnaceae* | H | N |  | + |  | 1 |  | + |
| *Lycopodium clavatum* L. | *Lycopodiaceae* | H | N |  |  |  | 1 |  | + |
| *Peperomia hernandifolia* (Vahl) A. Dietr. | *Piperaceae* | H | N |  |  |  |  | + |  |
| *Lycopodium cernuum* L. | *Lycopodiaceae* | H | N |  | 2 |  |  |  |  |
| *Odontosoria aculeata* (L.) J. Sm. | *Polypodiaceae* | H | N |  |  |  |  | + |  |
| *Isachne rigidifolia* (Poir.) Urb. | *Poaceae* | H | N |  |  |  |  |  | 1 |
| *Machaerina cubensis* (Kük.) T. Koyama | *Cyperaceae* | H | N |  |  |  |  |  | + |

Sites sampled. DR1.- Casabito. Ébano Verde (19340280E/2105321N). DR2.- Casabito (19340299E/2105967N). DR4.- Casabito. Ébano Verde (19340283N/2106095N). DR5.- Casabito. Ébano Verde (19340288E/2106283N). DR6.- Palmerito. Ébano Verde (19340165E/2106429N).
