## Supplemental file 2 for "The cloud forest in the Dominican Republic: diversity and conservation status"

**S2 Table 2.- *Ass.* *Cyatheo furfuracei-Prestoetum motanae***

|  | **Family** | **Biotype** | | **Status** |  |  |  |  |  |
| --- | --- | --- | --- | --- | --- | --- | --- | --- | --- |
| **Altitude** |  |  | |  | 1097 | 1373 | 1377 | 1251 | 1200 |
| **Area in m2 x 10** |  | |  |  | 200 | 50 | 100 | 100 | 50 |
| **Cover ratio In %** |  | |  |  | 100 | 100 | 100 | 100 | 100 |
| **Xn in m.** |  | |  |  | 20 | 9 | 9 | 15 | 7 |
| **Nº rel.** |  | |  |  | 6 | 13 | 14 | 15 | 17 |
| **Nº order** |  | |  |  | DR3 | DR7 | DR8 | DR9 | DR10 |
| **Characteristics of the association and higher units** | | | | | | | | | |
| *Prestoea montana* (Grah.) Nichol | *Arecaceae* | | A | N | 5 | 4 | 5 | 5 | 4 |
| *Arthrostylidium multispicatum* Pilger | *Poaceae* | | Tr | E | 2 | 3 | 2 | 1 | 2 |
| *Cyathea furfuracea* Baker | *Cyatheaceae* | | A | N | 2 | 1 | 2 | 2 | + |
| *Dendropanax arboreus* (L.) Dcne & Planch. | *Araliaceae* | | A | N | 2 |  | + | + | + |
| *3Alsophila minor* (D.C.Eaton) R.M.Tryon | *Cyatheaceae* | | A | N | 1 | 1 | 2 | 1 |  |
| *Ocotea leucoxylon* (Sw.) Mez | *Lauraceae* | | A | N |  | + | + | + | + |
| *Coccoloba wrightii* Lindau | *Polygnonaceae* | | A | N |  | 1 | + | 2 | + |
| *Alchornea latifolia* Sw. | *Euphorbiaceae* | | A | N | 2 | + |  | 1 |  |
| *Turpinia occidentalis* (Sw.) G. Don | *Staphyleaceae* | | A | N |  |  | + | 2 | 1 |
| *Brunellia comocladifolia* H. & B. | *Brunelliaceae* | | A | N | 2 |  |  |  | + |
| *Byrsonima lucida* (Mill.) L.c. Rich. | *Malpighiaceae* | | A | N |  | 1 |  |  | + |
| *Calyptrantes selleanus* Urb. & Ekm. | *Myrtaceae* | | A | E |  |  |  |  | + |
| *Cecropia screberiana* Miq. | *Moraceae* | | A | N | 2 |  | 2 |  |  |
| *Dichaea glauca* (Sw.) Lindley | *Orchidaceae* | | Ep | N |  | + |  | + | 1 |
| *Epidendrum anceps* Jacq. | *Orchidaceae* | | Ep | N |  | 1 |  |  |  |
| *Epidendrum jamaicense* Lindl | *Orchidaceae* | | Ep | N |  |  |  | + |  |
| *Epidendrum ramosum* Jacq. | *Orchidaceae* | | Ep | N |  |  | + | + |  |
| *Epidendrum ramosum* Jacq. | *Orchidaceae* | | Ep | N |  |  |  |  |  |
| *Grammitis asplenifolia* (L.) Proctor | *Grammitidaceae* | | Ep | N |  | + |  |  |  |
| *Guatteria blainii* (Griseb.) Urb. | *Annonaceae* | | A | N |  | + |  |  | + |
| *Guzmania monostrachya* (Sw.) Rusby | *Bromeliaceae* | | Ep | N |  | + | + |  |  |
| *Malpighia macracantha* Ekm. & Nied. | *Malpighiaceae* | | A | E |  |  |  | 2 |  |
| *Jacquiniella globosa* (Jacq.) Schlechter | *Orchidaceae* | | Ep | N |  | + |  |  |  |
| *Didymopanax tremulus* Krug. & Urb. | *Araliaceae* | | A | E | 1 |  |  |  |  |
| *Miconia mirabilis* (Aubl.) L.O. Willians | *Melastomataceae* | | A | N |  | + |  |  |  |
| *Exostema elliptica* Griseb. | *Rubiaceae* | | A | N |  |  | + |  |  |
| *Microgramma piloselloides* L. | *Polypodiaceae* | | Ep | N |  |  | + |  |  |
| *Camparettia falcata* Poepp. & Endl. | *Orchidaceae* | | Ep | N |  |  |  | + |  |
| *Antrophyum lanceolatum* (L.) Kaulf. | *Adiantaceae* | | Ep | N |  |  | + |  |  |
| *Myrsine coriacea* (Sw.) R. Br. | *Myrsinaceae* | | A | N |  | + |  |  | + |
| *Niphidium crassifolium* (L.) Lell. | *Polypodiaceae* | | Ep | N |  |  | + |  |  |
| *Oncidium variegatum* (Sw.) Sw. | *Orchidaceae* | | Ep | N |  |  |  | + |  |
| *Ophioglossum palmatum* L. | *Ophioglossaceae* | | Ep | N |  |  |  |  |  |
| *Phlebodium aureum* (L.) J. Smith | *Polypodiaceae* | | Ep | N |  |  |  | + |  |
| *Pleurothallis domingensis* Cogn. | *Orchidaceae* | | Ep | E |  | + |  |  | + |
| *Pothuya nudicaulis* (L.) Regel | *Bromeliaceae* | | Ep | N |  |  |  | + |  |
| *Rondeletia ochracea* Urb. | *Rubiaceae* | | A | E |  | + |  | 3 |  |
| **Companions species** |  | | | | | | | | |
| *Myrcia splendens*  (Sw.) DC. | *Myrtaceae* | | Ar | N |  | 5 | 2 | 2 | 5 |
| *Psychotria domingensis* Jacq. | *Rubiaceae* | | Ar | N |  | 3 | 3 | 1 | 1 |
| *Tabebuia bullata* A. Gentry | *Bignoniaceae* | | Ar | E | 1 |  | + | + | + |
| *Blechnum tuerckheimii* A. Brause | *Blechnaceae* | | H | E |  | 1 | 2 | 3 |  |
| *Psychotria guadalupensis* (DC.) Howard | *Rubiaceae* | | Ar | N |  | 3 |  | 1 | 1 |
| *Renealmia jamaicensis* (Gaertn.) Horan var. *puberula* (Gagn.) Maas | *Zingiberaceae* | | H | N |  | 2 | 2 |  | + |
| *Mikania venosa* A. Liogier | *Asteraceae* | | Tr | E |  |  | + | + | 2 |
| *Sagraea fuertesii* (Cogn.in Urb.) Alain | *Melastomataceae* | | Ar | E |  | 1 |  |  | 1 |
| *Senecio lucens* (Poir) Urb. | *Asteraceae* | | Tr | E |  |  | + | 2 | 1 |
| *Smilax havanensis* Jacq. | *Smilacaceae* | | Tr | N |  | + |  |  |  |
| *Solanum crotonoides* Lam. | *Solanaceae* | | Ar | N |  | 1 |  |  | + |
| *Solanum virgatum* Lam. | *Solanaceae* | | Ar | N |  |  | + |  |  |
| *Stigmaphyllon emarginatum* (L.) A. Juss. | *Malpighiaceae* | | Tr | N |  |  |  |  | + |
| *Uncinia hamata* (L.) Urb. | *Cyperaceae* | | H | N |  | + | + | + |  |
| *Vaccinium racemosum* (Vahl) Wilbur & Luteyn | *Ericaceae* | | Tr | N |  | + |  |  |  |
| *Vitis tiliifolia* H. & B. ex Willd. | *Vitaceae* | | Tr | N |  |  |  |  | + |
| *Vittaria lineata* (L.) Smith | *Pteridaceae* | | Ep | N |  |  | + |  |  |
| *Blechnum occidentale* L. | *Blechnaceae* | | H | N |  | 1 | 2 |  | + |
| *Cestrum coelophlebium* O. E. Schulz | *Solanaceae* | | Ar | E |  |  |  | 1 | + |
| *Cestrum inclusum* Urb. | *Solanaceae* | | Ar | E |  |  | 5 |  |  |
| *Cissampelos pareira* L. | *Menispermiaceae* | | Tr | N | 1 |  | + |  |  |
| *Commelina elegans* Kunth | *Commelinaceae* | | H | N |  |  | + |  |  |
| *Daphnosis crassifolia* (Poir.) Meiss. | *Thymelaeaceae* | | Ar | N |  | + |  |  |  |
| *Diplazium hastile* (Christ.) C. Chr. | *Athyriaceae* | | H | N |  |  | 2 |  |  |
| *Diplazium hians* Kuntze | *Athyriaceae* | | H | N |  |  |  | 2 |  |
| *Gleychenia bifida* (Willd.) Spreng. | *Gleycheniaceae* | | H | N | 1 |  |  |  | + |
| *Gomedesia lindeniana* Berg. | *Myrtaceae* | | Ar | N |  |  |  |  | 1 |
| *Gyrotaenia myriocarpa* Griseb. | *Urticaceae* | | Ar | N |  |  | + |  |  |
| *Hyptis americana* (Poir.) Briq. | *Lamiaceae* | | Ar | N |  | + |  |  |  |
| *Ichnanthus pallens* (Sw.) Munro | *Poaceae* | | H | N |  | 1 | + | + |  |
| *Ipomoea furcyensis* Urb. | *Convolvulaceae* | | Tr | E |  |  |  | + |  |
| *Lasianthus lanceolatus* (Griseb.) Gómez Maza | *Rubiaceae* | | Ar | N |  | 1 |  |  |  |
| *Lobelia robusta* Graham | *Campanulaceae* | | Ar | E |  |  | + |  |  |
| *Lobelia rotundifolia* Juss. | *Campanulaceae* | | H | E |  | + |  |  |  |
| *Odontadenia polyneura* (urb.) Wood. | *Apocynaceae* | | Tr | E |  |  |  |  | 1 |
| *Odontosoria uncinella* (Kunze) Fée | *Polypodiaceae* | | Tr | N |  | + |  |  |  |
| *Olyra latifolia* L. | *Poaceae* | | H | N |  |  |  | + |  |
| *Palicourea alpina* (Sw.) DC. | *Rubiaceae* | | Ar | N |  | + |  |  | + |
| *Peperomia hernandifolia* (Vahl) A. Dietr. | *Piperaceae* | | H | N |  | + |  |  |  |
| *Pilea geminata* Urb. | *Urticaceae* | | H | E |  |  | 2 |  |  |
| *Polypodium loriceum* L. | *Polypodiaceae* | | Ep | N |  |  |  | + |  |
| *Pothomorphe peltata* (L.) Miquel | *Piperaceae* | | Ar | N |  |  | + |  |  |
| *Mucuna urens* (L.) Fawc. & Rendle | *Fabaceae* | | Tr | N |  | + |  |  |  |
| *Myrcia deflexa* (Poir) DC. | *Myrtaceae* | | Ar | N |  |  |  | 1 | + |

Sites sampled. DR3.- Río Jatubei (19341984E/2105891N). DR7.- Camino Casabito al Arroyazo (10339971E/2105962N). DR8.- Bajada Casabito al Centro Fernándo Dominguez (19339590E/2105699N). DR9.- Casabito-Arroyazo (Ébano Verde) (19339203E/2105784N). DR10.- Near Arroyazo (19339203E/2105785N).
