## Supplemental file 3 for "The cloud forest in the Dominican Republic: diversity and conservation status"

**S3 Table 3. *Ass. Hyeronimo dominguensis-Magnolietum hamorii*.**

|  | **Family** | **Biotype** | **Status** |  |  | | |  | |
| --- | --- | --- | --- | --- | --- | --- | --- | --- | --- |
| **Altitude** |  |  |  | 1207 | | 1239 | 1233 | | 1140 |
| **Area in m2 x 10** |  |  |  | 200 | | 200 | 200 | | 200 |
| **Cover ratio In %** |  |  |  | 100 | | 100 | 100 | | 100 |
| **Xn in m.** |  |  |  | 25 | | 15 | 20 | | 15 |
| **Nº rel.** |  |  |  | 23 | | 24 | 25 | | 26 |
| **Nº order** |  |  |  | DR11 | | DR12 | DR13 | | DR14 |
| **Characteristics of the association and higher units** |  |  |  |  | |  |  | |  |
| *Magnolia hamorii* Howard | *Magnoliaceae* | A | E | 5 | | 2 | 2 | | 5 |
| *Hyeronima domingensis* Urb. | *Euphorbiaceae* | A | E | 5 | | 2 | 5 | | + |
| *Cyathea fulgens* C. Chr. | *Cyatheaceae* | A | N | 2 | | 2 | 2 | | 1 |
| *Myrsine coriacea* (Sw.) R. Br. | *Myrsinaceae* | A | N | 1 | | 1 | 1 | | 2 |
| *Didymopanax tremulus* Krug. & Urb. | *Araliaceae* | A | E | + | | 5 | 2 | | 3 |
| *Brunellia comocladifolia* H. & B. | *Brunelliaceae* | A | N | 2 | | 1 |  | |  |
| *Prestoea montana*(Grah.) Nichol | *Arecaceae* | A | N | + | | 2 | 2 | | 3 |
| *Beilschmiedia pendula* (Sw.) Hemsl. | *Lauraceae* | A | N | 2 | |  | 1 | |  |
| *Ocotea leucoxylon* (Sw.) Mez | *Lauraceae* | A | N |  | | 1 | 1 | | + |
| *Calyptrantes selleanus* Urb. & Ekm. | *Myrtaceae* | A | E | + | | 1 | 1 | |  |
| *Weinmannia pinnata* L. | *Cunoniaceae* | A | N | 2 | | 2 | 2 | |  |
| *Pleurothalis ruscifolia* (Jaq.) R. Br. | *Orchidaceae* | Ep | N | 1 | | 2 | 2 | |  |
| *Elleanthus cephalotus* Garay & Sweet | *Orchidaceae* | Ep | N | 2 | | 2 | 1 | |  |
| *Elaphoglossum crinitum* (L.) C. Chr. | *Lomariopsidaceae* | Ep | N | 1 | | 1 | + | |  |
| *Columnea sanguinea* Urb. | *Gesneriaceae* | ArEp | N | 1 | | 2 | 1 | |  |
| *Elaphoglossum latifolium* (Sw.) J. Sm. | *Lomariopsidaceae* | Ep | N | 2 | | 2 | 2 | |  |
| *Miconia prasina* (Sw.) DC. | *Melastomataceae* | A | N | 1 | |  | 1 | |  |
| *Rondeletia ochracea* Urb. | *Rubiaceae* | A | E | 1 | | 1 | 1 | |  |
| *Alchornea latifolia* Sw. | *Euphorbiaceae* | A | N | 1 | |  |  | | + |
| *Dendropanax arboreus* (L.) Dcne & Planch. | *Araliaceae* | A | N | 1 | |  |  | |  |
| *Miconia mirabilis* (Aubl.) L.O. Willians | *Melastomataceae* | A | N |  | | 1 |  | | + |
| *Epidendrum ramosum* Jacq. | *Orchidaceae* | Ep | N |  | |  | 2 | | + |
| *Ophioglossum palmatum* L. | *Ophioglossaceae* | Ep | N | + | | 1 |  | |  |
| *Ocotea acarina* C.K. Allen | *Lauraceae* | A | E |  | |  | 2 | | 1 |
| *Chionanthus domingensis* Lam. | *Oleaceae* | A | N |  | |  | 2 | |  |
| *Ocotea nemodaphne* Mez | *Lauraceae* | A | N |  | | 1 |  | |  |
| *Ilex macfadyenii* (Walp.) Rehder | *Aquifoliaceae* | A | N |  | | 1 |  | |  |
| *Niphidium crassifolium* (L.) Lell. | *Polypodiaceae* | Ep | N | 2 | |  |  | |  |
| *Polypodium loriceum* L. | *Polypodiaceae* | Ep | N | 1 | |  |  | |  |
| *Epidendrum jamaicense* Lindl | *Orchidaceae* | Ep | N |  | | 2 |  | |  |
| *Phlebodium aureum* (L.) J. Smith | *Polypodiaceae* | Ep | N | 1 | |  |  | |  |
| *Dichaea glauca* (Sw.) Lindley | *Orchidaceae* | Ep | N |  | | 2 |  | |  |
| *Epidendrum carpophorum* Barb. Rodr. | *Orchidaceae* | Ep. | N |  | |  | 1 | |  |
| *Ocotea floribunda* (Sw.) Mez | *Lauraceae* | A | N |  | |  |  | | 1 |
| *Anacheilium cochleatum* (L.) Hoffm. | *Orchidaceae* | Ep | N |  | |  |  | | + |
| *Ocotea patens* (Sw.) Nees | *Lauraceae* | A | N |  | |  |  | | + |
| *Guarea guidonea* Sleumer | *Meliaceae* | A | N | 1 | |  |  | |  |
| *Maxillaria coccinea* (Jacq.) L.O. Wms. | *Orchidaceae* | Ep | N |  | | 2 |  | |  |
| *Ocotea foeniculacea* Mez | *Lauraceae* | A | N |  | | 1 |  | |  |
| *Cecropia screberiana* Miq. | *Moraceae* | A | N |  | |  |  | | 1 |
| *Beilschmiedia pendula* (Sw.) Hemsl. | *Lauraceae* | A | N |  | |  |  | | 1 |
| **Companions species** |  |  |  |  | |  |  | |  |
| *Psychotria domingensis* Jacq. | *Rubiaceae* | Ar | N | 2 | | 2 | 2 | | 1 |
| *Mikania venosa A. Liogier* | *Asteraceae* | Tr | E | 1 | | 2 | 1 | | 2 |
| *Gomedesia lindeniana* Berg. | *Myrtaceae* | Ar | N | 1 | | 1 | 1 | | 2 |
| *Lasianthus bahorucanus* Zanoni | *Rubiaceae* | H | E | 2 | | 2 | 1 | | 1 |
| *Columnea domingensis* (Urb.) Wiehler | *Gesneriaceae* | Ar | E | 2 | | 1 | + | | 1 |
| *Odontosoria uncinella* (Kunze) Fée | *Polypodiaceae* | Tr | N | 3 | | 2 | 2 | | 2 |
| *Mecranium ovatum* Cog. | *Melastomataceae* | Ar | E | 2 | | 1 | 1 | | 1 |
| *Vriesea tuercheimii* (Mez.) L.B. Smith | *Bromeliaceae* | H | E | 2 | | 2 | 2 | | 1 |
| *Nephrolepis biserrata* (Sw.) Schott | *Lomariopsidaceae* | H | N | 2 | | 2 | 2 | | 2 |
| *Peperomia hernandifolia* (Vahl) A. Dietr. | *Piperaceae* | H | N | + | | 1 | 1 | | 1 |
| *Psychotria guadalupensis* (DC.) Howard | *Rubiaceae* | Ar | N | 2 | | 2 | 1 | |  |
| *Myrcia deflexa* (Poir) DC. | *Myrtaceae* | Ar | N | 2 | | 1 | 1 | | 2 |
| *Lomariposis sorbifolia* (L.) Feé | *Lomariopsidaceae* | H | N | 1 | |  | 1 | | 1 |
| *Hedyosmum domingense* Urb. | *Chloranthaceae* | Ar | E |  | | 1 | 1 | | + |
| *Lomariposis sorbifolia* (L.) Feé | *Lomariopsidaceae* | H | N | 2 | | 2 |  | | 1 |
| *Renealmia jamaicensis* (Gaertn.) Horan var. *puberula* (Gagn.) Maas | *Zingiberaceae* | H | N | 2 | | 1 | 2 | |  |
| *Vaccinium racemosum* (Vahl) Wilbur & Luteyn | *Ericaceae* | Tr | N |  | | 1 | 1 | |  |
| *Macrocarpaea domingensis* Urb. | *Gentianaceae* | Ar | E |  | | 2 | 1 | |  |
| *Polygala fuertesii* (Urb.) Blake | *Polygalaceae* | Ar | E |  | | 1 | 1 | |  |
| *Arthrostylidium multispicatum* Pilger | *Poaceae* | Tr | E | 3 | | 2 |  | |  |
| *Torralbasia cuneifolia* (C. Wright) Krug. & Urb. | *Celastraceae* | Ar | N |  | | 1 | 1 | |  |
| *Mucuna urens* (L.) Fawc. & Rendle | *Fabaceae* | Tr | N | 1 | |  |  | | 2 |
| *Schlegelia brachyantha* Griseb. | *Schlegeliaceae* | Tr | N | 1 | |  |  | | + |
| *Meriania involucrata* (Desv.) Naud. | *Melastomataceae* | Ar | E |  | | 1 | 1 | |  |
| *Hypolepis hispaniolica* Mason | *Polypodiaceae* | Tr | E |  | | 2 |  | | 1 |
| *Arthrostylidium sarmentosum* Pilger | *Poaceae* | Tr | N |  | | 2 | 2 | |  |
| *Blechneum fragile* (Liebm.) Morton & Lellinger | *Blechnaceae* | H | N |  | | 2 | 2 | |  |
| *Ilex tuerckheimii* Loes. | *Aquifoliaceae* | Ar | E |  | |  | + | |  |
| *Cordia dependens* Urb. & Ekm. | *Boraginaceae* | Ar | E |  | |  |  | | + |
| *Passiflora rubra* L. | *Passifloraceae* | Tr | N |  | |  |  | | + |
| *Eupatorium odoratum* L. | *Asteraceae* | Ar | N |  | |  |  | | + |
| *Mikania cordifolia* (L.) Willd. | *Asteraceae* | Tr | N |  | |  |  | | 1 |
| *Psychotria liogieri S*ateyerm | *Rubiaceae* | Ar | N |  | |  |  | | + |
| *Marattia kaulfussii* J. Smith | *Marattiaceae* | H | N | 1 | |  |  | |  |
| *Asplenium radicans* L. | *Aspleniaceae* | H | N | 1 | |  |  | |  |
| *Smilax domingensis* Willd. | *Smilacaceae* | Tr | N | + | |  |  | |  |
| *Leandra limoides* (Urb.) W. Judd & Skean | *Melastomataceae* | Ar | E |  | | 1 |  | |  |
| *Hillia parasitica* Jacq. | *Rubiaceae* | Tr | N |  | | 2 |  | |  |
| *Cestrum daphnoides* Griseb. | *Solanaceae* | Ar | E |  | | 1 |  | |  |
| *Tibouchina longifolia* (Vahl) Baill. | *Melastomataceae* | Ar | N |  | | 1 |  | |  |
| *Clidemia umbellata* (Miller) L.O. Wms. | *Melastomataceae* | Ar | N |  | |  |  | | + |
| *Schradera subsessilis* Steyermark | *Rubiaceae* | Tr | E | 1 | |  |  | |  |
| *Marcgravia rubra* A. Liogier | *Marcgraviaceae* | Tr | E |  | |  | 1 | |  |
| *Lobelia rotundifolia* Juss. | *Campanulaceae* | H | E |  | | 1 |  | |  |
| *Blechnum occidentale* L. | *Blechnaceae* | H | N |  | |  |  | | + |
| *Cissampelos pareira* L. | *Menispermiaceae* | Tr | N |  | |  |  | | + |
| *Myrcia splendens*  (Sw.) DC. | *Myrtaceae* | Ar | N |  | |  |  | | 3 |
| *Ichnanthus pallens* (Sw.) Munro | *Poaceae* | H | N |  | |  |  | | 1 |
| *Sagraea fuertesii* (Cogn.in Urb.) Alain | *Melastomataceae* | Ar | E |  | | 1 |  | |  |

Sites sampled. DR11.- Sierra Bahoruco. El Cachote (19267592E/2002124N). DR12.- Sierra Bahoruco. El Cachote (19268161E/2002764N). DR13.- Sierra Bahoruco. Prox. el Cachote (19268152E/2002964N). DR14.- Km. 3 del poblado Cachote (19268736E/2000217N).
