## Supplemental file 4 for "The cloud forest in the Dominican Republic: diversity and conservation status"

**S4 Table 4.- *Ass. Ormosio krugii-Prestoetum montanae*.**

|  | **Family** | **Biotype** | **Status** |  |  | | |
| --- | --- | --- | --- | --- | --- | --- | --- |
| **Altitude** |  |  |  | 519 | | 541 | 530 |
| **Area in m2 x 10** |  |  |  | 200 | | 200 | 200 |
| **Cover ratio In %** |  |  |  | 75 | | 100 | 100 |
| **Xn in m.** |  |  |  | 15 | | 12 | 15 |
| **Nº rel.** |  |  |  | 13 | | 15 | 15b |
| **Nº order** |  |  |  | DR15 | | DR16 | DR17 |
| **Characteristics of the association and higher units** |  |  |  |  | |  |  |
| *Prestoea montana* (Grah.) Nichol | *Arecaceae* | A | N | 3 | | 4 | 4 |
| *Cecropia screberiana* Miq. | *Moraceae* | A | N | 3 | | 2 | 3 |
| *Alchornea latifolia* Sw. | *Euphorbiaceae* | A | N | 2 | | 5 | 4 |
| *Miconia mirabilis* (Aubl.) L.O. Willians | *Melastomataceae* | A | N | 3 | | 2 | 2 |
| *Miconia prasina* (Sw.) DC. | *Melastomataceae* | A | N | 1 | | 1 | 1 |
| *Guarea guidonea* Sleumer | *Meliaceae* | A | N | + | | 4 | 4 |
| *Cyathea arborea* (L.) J.E. Smith | *Cyatheaceae* | A | N | 3 | | 4 | 4 |
| *Turpinia occidentalis* (Sw.) G. Don | *Staphyleaceae* | A | N | 1 | | + | 1 |
| *Clusia rosea* Jacq. | *Clusiaceae* | A | N | 1 | | + | + |
| *Ocotea globosa* (Aubl.) Schlecht. & Cham. | *Lauraceae* | A | N | 2 | | 1 | 1 |
| *Casearea arborea* (L.C.Rich.) Urb. | *Flacourtiaceae* | A | N | 1 | | 1 | + |
| *Oreopanax capitatus* (Jacq.) Decne. & Planch. | *Araliaceae* | A | N | 2 | | 3 | 3 |
| *Didymopanax morototoni* (Aubl.) Decne. & Planch | *Araliaceae* | A | N | 2 | | 3 | 3 |
| *Byrsonima spicata* (Cav.) Kunth | *Malpighiaceae* | A | N | + | | 1 | 1 |
| *Buchenavia tetraphylla* (Aubl.) R. A. Howard | *Combretaceae* | A | N | 1 | | 1 | 1 |
| *Sloanea berteriana* Choisy | *Elaeocarpaceae* | A | N | 1 | | 1 | 2 |
| *Ormosia krugii* Urb. | *Fabaceae* | A | N | 2 | | 2 | 2 |
| *Miconia serrulata* (DC.) Naud. | *Melastomataceae* | A | N | + | | + | 1 |
| *Bactris plumeriana* Mart. | *Arecaceae* | A | E |  | | 1 | 1 |
| *Myrsine coriacea* (Sw.) R. Br. | *Myrsinaceae* | A | N | 1 | | 1 |  |
| *Ocotea leucoxylon* (Sw.) Mez | *Lauraceae* | A | N |  | | 2 | 2 |
| *Inga fagifolia* (L.) Willd. ex Benth. | *Mimosaceae* | A | N |  | | + | + |
| *Inga vera* Willd. | *Mimosaceae* | A | N |  | | + | + |
| *Cupania americana* L. | *Sapindaceae* | A | N | 2 | |  |  |
| *Hirtella triandra* Sw. | *Chrysobalanaceae* | A | N | + | |  |  |
| *Miconia racemosa* (Aubl.) DC. | *Melastomataceae* | A | N | 1 | |  |  |
| *Zantoxylum martinicensis* (Lam.) DC. | *Rutaceae* | A | N | 1 | |  |  |
| *Guzmania monostrachya* (Sw.) Rusby | *Bromeliaceae* | Ep | N | + | |  |  |
| *Microgramma piloselloides* L. | *Polypodiaceae* | Ep | N | + | |  |  |
| **Companions species** |  |  |  |  | |  |  |
| *Cnemidaria horrida* (L.) K. Presl | *Cyatheaceae* | Ar | N | 2 | | 2 | 2 |
| *Cnemidaria horrida* (L.) K. Presl | *Cyatheaceae* | Ar | N | 2 | | 2 | 2 |
| *Pytirogramma calomelanos* (L. ) Link | *Polypodiaceae* | H | N | 1 | | + | + |
| *Ipomoea tiliacea* (Willd.) Choisy | *Convolvulaceae* | Tr | N | + | | 2 | 2 |
| *Mucuna urens* (L.) Fawc. & Rendle | *Fabaceae* | Tr | N | 1 | | 2 | 2 |
| *Solanum torvum* Sw. | *Solanaceae* | Ar | N | 1 | | 1 | 1 |
| *Mikania cordifolia* (L.) Willd. | *Asteraceae* | Tr | N | 2 | | 1 | 1 |
| *Psychotria domingensis* Jacq. | *Rubiaceae* | Ar | N |  | | 2 | 1 |
| *Pothomorphe peltata* (L.) Miquel | *Piperaceae* | Ar | N |  | | 2 | 2 |
| *Tibouchina longifolia* (Vahl) Baill. | *Melastomataceae* | Ar | N | 1 | |  | + |
| *Nepsera aquatica* (Aubl.) Naud. | *Melastomataceae* | Ar | N | 1 | |  | + |
| *Syngonium podophyllum* Schott | *Araceae* | Tr | N | 2 | | + |  |
| *Urera baccifera* (L.) Gaud. | *Urticaceae* | Ar | N |  | | 2 | 2 |
| *Psychotria uliginosa* Sw. | *Rubiaceae* | Ar | N |  | | 2 | 2 |
| *Coccocypselum herbaceum* Aubl. | *Rubiaceae* | H | N | + | |  |  |
| *Piper adunculum* L. | *Piperaceae* | Ar | N | 1 | |  |  |
| *Cissus verticillata* (L.) Nicholson & Farris | *Vitaceae* | Tr | N | 1 | |  |  |
| *Neurolaena lobata* (L.) Cass. | *Asteraceae* | H | N | + | |  |  |
| *Triunfetta semitriloba* Jacq. | *Tiliaceae* | H | N | 1 | |  |  |
| *Clidemia umbellata* (Miller) L.O. Wms. | *Melastomataceae* | Ar | E | 1 | |  |  |
| *Gleychenia bifida* (Willd.) Spreng. | *Gleycheniaceae* | H | N | 1 | |  |  |
| *Lycopodium clavatum* L. | *Lycopodiaceae* | H | N | 1 | |  |  |
| *Ichnanthus pallens* (Sw.) Munro | *Poaceae* | H | N | 1 | |  |  |
| *Nephrolepis multiflora* (Roxb.) Jarret | *Lomariopsidaceae* | H | N | 1 | |  |  |
| *Smilax domingensis* Willd. | *Smilacaceae* | Tr | N | + | |  |  |
| *Mimosa pudica* L. | *Mimosaceae* | H | N | 1 | |  |  |
| *Palicourea crocea* (Sw.) Schultes | *Rubiaceae* | Ar | N | 1 | |  |  |
| *Urena lobata* L. | *Malvaceae* | Ar | N | 1 | |  |  |
| *Hedychium coronarium* Koen. | *Zingiberaceae* | H | I | 1 | |  |  |
| *Solanum jamaicense* Mill. | *Solanaceae* | Ar | N | 1 | |  |  |
| *Entada gigas* (L.) Fawc. & Rendle | *Fabaceae* | Tr | N | + | |  |  |

Sites sampled: DR15.- El Trece (eastern range) (19Q0489524/2092418). DR16.- Dieciseis de Mitche (19Q0486735/2092513). DR17.- Near Dieciseis de Mitche (19Q0486736/2092514).
